# Model-based prediction of bacterial population dynamics in gastrointestinal infection

**DOI:** 10.1101/2020.08.11.244202

**Authors:** Janina K. Geißert, Erwin Bohn, Reihaneh Mostolizadeh, Andreas Dräger, Ingo B. Autenrieth, Sina Beier, Oliver Deusch, Martin Eichner, Monika S. Schütz

## Abstract

The complex interplay of a pathogen with the host immune response and the endogenous microbiome determines the course and outcome of gastrointestinal infection (GI). Expansion of a pathogen within the gastrointestinal tract implies an increased risk to develop systemic infection. Through computational modeling, we aimed to calculate bacterial population dynamics in GI in order to predict infection course and outcome. For the implementation and parameterization of the model, oral mouse infection experiments with *Yersinia enterocolitica* were used. Our model takes into account pathogen specific characteristics, such as virulence, as well as host properties, such as microbial colonization resistance or immune responses. We were able to confirm the model calculations in these scenarios by experimental mouse infections and show that it is possible to computationally predict the infection course. Far future clinical application of computational modeling of infections may pave the way for personalized treatment and prevention strategies of GI.

## Introduction

Gastrointestinal infection is a frequent disease that causes significant morbidity and economic burden (Dautzenberg et al., 2015; Jia et al., 2019). Being self-resolving in most cases, symptomatic treatment (e.g. rehydration) is sufficient for otherwise healthy individuals.

In contrast, gastrointestinal tract (GIT) infection can cause high morbidity and even fatal disease in healthcare settings and in specific populations such as newborns, elderly and immunocompromised individuals. According to the OECD Health Report, 2016-2017, ∼ 9 % of healthcare-associated infections were related to the GIT (OECD/European Union Paris/European Union, (2018)). At present it is not possible to reliably identify patients who are at risk of developing a fatal systemic disease, and we do not have personalized prevention strategies. Thus, it would be desirable to develop a means to identify high-risk individuals and to use this knowledge to stratify patient treatment.

In recent years the benefit of computational methods to improve patient treatment has been recognized and we know that the efficacy of drug treatment is highly variable between individuals. Numerous host factors, such as the composition of the microbiota, can significantly influence the success of a treatment (Guthrie & Kelly, 2019). Therefore, computational approaches are being developed that integrate available patient data to derive personalized and improved therapy guidelines (Shameer et al., 2015; Toh et al., 2019). Related to such approaches, we asked whether we could use computational modeling to predict the population dynamics of an enteropathogen within the GIT, and thereby predict the infection course and its likely outcome. With the ability to integrate the host and pathogen specific properties that the most influence the course and outcome of GIT infection (i.e., virulence factors expressed by the pathogen, presence of a microbiota, immune competence), such a model could be a helpful tool to identify individuals at particularly increased risk of developing a fatal disease.

To tackle our question, we chose a mouse model of infection that makes the abovementioned factors accessible and modifiable. Experimental mouse infections were used to generate a dataset to build up, parametrize, and evaluate the model. *Yersinia enterocolitica* (Ye) was employed as a model enteropathogen because it has well studied virulence factors. The most important ones are the adhesin YadA, which mediates attachment to host cells (El Tahir & Skurnik, 2001), and the type three secretion system (T3SS), which mediates immune evasion (Cornelis, 2002; Ruckdeschel et al., 1996). Both contribute to the efficient colonization of the intestinal tract which elicits an inflammatory response that leads to a reduction of density and complexity of the commensal microbiota (Lupp et al., 2007; Stecher et al., 2007). Strains deficient for either YadA or a functional T3SS were used to modulate the virulence of Ye and to find out whether these traits affect the infection course. To mimic host microbiota deficiency, we used germfree (GF) mice. As model for an immune-compromised host, we used *MyD88*^-/-^ mice, which are strongly impaired in their antimicrobial immune responses.

Our study provides proof-of-concept that it is possible to create a computational model of gastrointestinal infection and underlines the validity of such approaches. To create our model, a reasonable knowledge about the infection biology of the causative pathogen was essential. In addition, the accurate definition and determination of parameter values were crucial. In the future, sophisticated computational models could be developed and applied in clinical routine to identify high-risk patients and to stratify their treatment in function of the specific properties of the individual patient and the causative pathogen. Moreover, such models will contribute to stimulate new hypotheses and provide novel mechanistic insights into the course of gastrointestinal infections.

## Results

The final aim of this study was to create a computational model that can predict the course of a GIT infection by calculating the colony forming units (CFU) in feces as a surrogate. Our model development process consisted of 6 steps: **Step 1** was the creation of a primary experimental dataset for Ye population dynamics in a host harboring a diverse microbiota and an intact immune response, i.e. C57BL/6J wild type mice with a specific pathogen-free (SPF) microbiota. In **step 2**, based on the data from step 1, we generated hypotheses about Ye population dynamics in the absence of microbiota and in an immunocompromised host. **Step 3** was to define the most critical interactions between Ye, the host immune system, and the microbiota. **Step 4** was the mathematical description of Ye population dynamics, and **step 5** included the experimental determination of specific parameter values and the calibration for parameters with unknown values. Finally, in s**tep 6**, we evaluated the model output by comparison to the experimental data that were obtained by infections of immunocompetent SPF wild type mice, of a host lacking microbiota (C57BL/6J germfree (GF) mice), and of an immunocompromised host with a diverse microbiota (C57BL/6J *MyD88*^-/-^ SPF mice).

(**Step 1**): C57BL/6J wild type SPF mice were infected with a 1:1 mixture of the Ye wild type (wt) strain and either the Ye YadA0 mutant strain, which lacks the adhesin YadA, or the Ye T3S0 mutant strain, which is impaired in type three secretion. We then determined the bacterial counts of Ye wt and the co-infected mutant strains in feces by serial plating on selective media (**Fig. 1A, C**). We found that the Ye wt strain was able to stably colonize the GIT of all animals over the entire observation period of 14 days. In contrast, the bacterial counts of both the Ye YadA0 and the T3S0 mutant strains never reached Ye wt levels and dropped below our limit of detection on 10 dpi. The competitive indices show the reduced virulence of Ye YadA0 and Ye T3S0 compared to wild type (**Fig. 1B, D**). We recorded the individual body weight of animals, as a sustainable weight loss is a sign of severe infection and fatal outcome. We found that three (out of 14) animals in the Ye wt : Ye YadA0 and four in the Ye wt : Ye T3S0 coinfection group significantly lost weight from 3 dpi on, while the mean change in body weight of all other mice slightly increased or remained static (**Fig. S1A and B;** suppl. files can be found after the references section). The mean gain of weight of uninfected animals over a time-course of 14 days was ∼7 g during previous studies. Infected animals that were not affected by weight-loss gained weight at a comparable level. Thus, we do not assume that those animals were suffering from severe infection. The most striking difference between Ye wt : Ye YadA0 and Ye wt : Ye T3S0 coinfections was that the bacterial counts of the Ye T3S0 strain peaked later and at considerably lower levels compared to both Ye wt, and the Ye YadA0 mutant strain.

**Figure 1.**
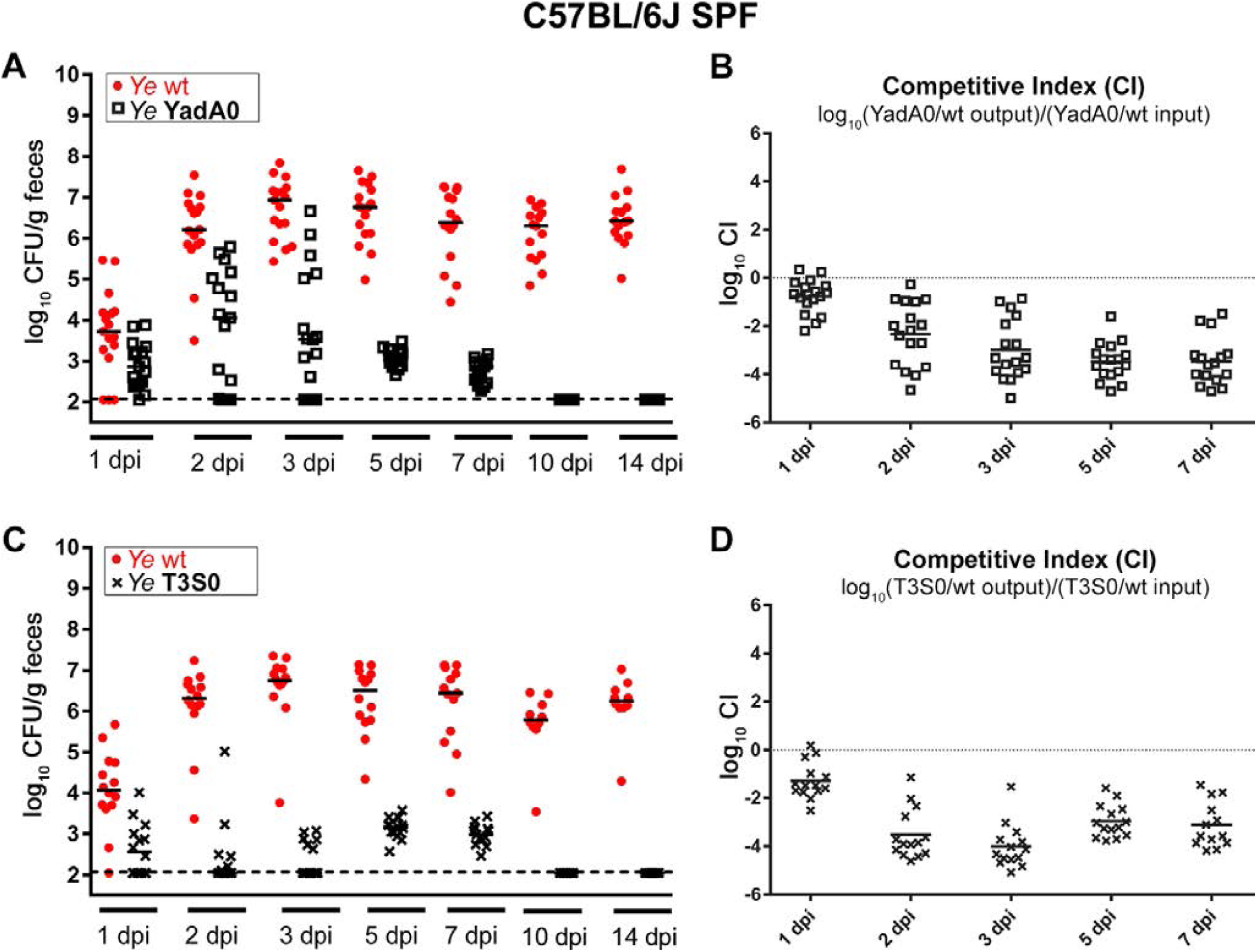
Ye population dynamics during coinfection of SPF-colonized mice. **(A)** Colony forming units (CFU) in feces of individual animals (n = 14) and the median thereof after oral 1:1 coinfection of C57BL/6J SPF mice with a Ye wild type (wt) strain and an attenuated mutant strain lacking the *Yersinia* adhesin A (Ye YadA0). The limit of detection is indicated by a dashed line. **(B)** The competitive index (CI) of the Ye wt : Ye YadA0 coinfection was calculated as indicated. A negative CI is indicative of an attenuation of the mutant strain. **(C)** CFU in feces of individual mice after coinfection with Ye wt and a mutant being impaired in type three secretion (Ye T3S0). **(D)** CI of the Ye at Ye wt : Ye T3S0 coinfection.

In summary, these data indicate that the pleiotropic functions of YadA and the effector functions mediated by the T3SS seem to be crucial for effective immune evasion and colonization of the GIT in the presence of a complex microbiome and an immunocompetent host. This effect has been shown in coinfection for the first time but has been demonstrated previously in oral single-infections using the YadA deficient strain and in coinfections with a strain lacking the single effector protein Yop H (Dave et al., 2016; Di Genaro et al., 2003). The body weight development indicates that, at later time points of infection, a kind of balanced state might be reached again, although Ye wt still colonizes the GIT at high CFUs.

In **step 2**, based on the results of step 1, we inferred Ye population dynamics in a host lacking microbiota and in an immunocompromised host (**Fig. 2**). We know that in SPF wild type animals (**Fig. 2A and B)** Ye elicits an innate immune response, leading to an increased antimicrobial peptide (AMP) and cytokine production as well as infiltration of professional phagocytes into the mucosal site (Handley et al., 2004; Pepe et al., 1995) (**Fig. S2**). This immune response more strikingly affects the endogenous microbiota and reduces its density and complexity (**Fig. S3**), especially in locations close to the epithelium. In the following, we refer to these locations as the mucosal compartment, comprising the mucosa, the epithelium, and the gut-associated lymphatic tissues, such as the Peyers Patches (PP) and the overlying microfold cells (M-cells).

**Figure 2.**
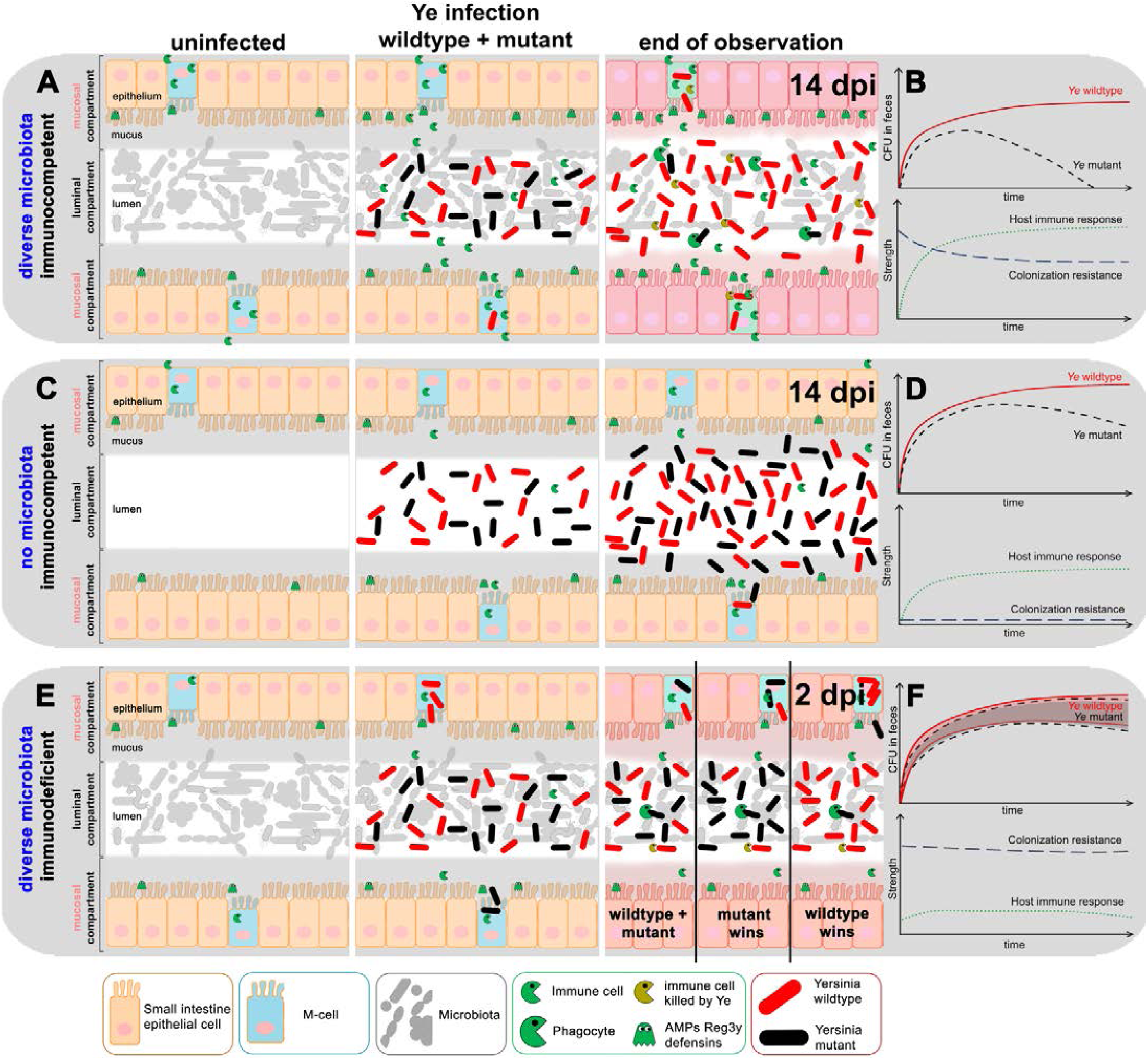
Schematic overview of the presumed infection progression after coinfection of different mouse models with Ye wt and mutant strains. **(A)** Scheme of the small intestine of SPF-colonized C57BL/6J wildtype mice during homeostasis (left), after initial disturbance (mid) and expected outcome after coinfection with a 1:1 mixture of Ye wt and an attenuated mutant strain. Initially, the gut lumen in SPF mice is densely colonized with a complex microbiota. Ye infection, associated with an infiltration of microfold cells (M-cells) mainly conducted by the wt strain, leads to an unspecific antimicrobial immune response accompanied by the release of phagocytic cells into the gut lumen and augmented expression of antimicrobial proteins (AMPs, Reg3γ, defensins) by epithelial cells. Both the antimicrobial response and inflammation affect at least parts of the microbiota and reduce its complexity and density. Whereas Ye wt can counteract phagocytosis by injection of effectors into immune cells and thereby kills them, the Ye mutant strain is more susceptible to phagocytosis and killing by immune cells and thus is finally outcompeted 14 days after infection onset. (**B**) Schematic overview of expected Ye wt and mutant CFU in feces during the infection course (upper diagram) and the presumed strength of host immune response and colonization resistance (CR; bottom diagram). **(C)** In germ-free (GF) mice that lack a microbiota that confers CR and harbor an immature immune system, Ye wt, and mutant strain are both able to colonize the gut lumen and do not necessarily need to enter a mucosa-near site to colonize the gut effectively. This leads only to weak antimicrobial responses that Ye can cope with, without the necessity to possess specific virulence traits (such as YadA or a functional T3SS). This results in comparable numbers of wt and mutant strains at the end of the observation period. **(D)** Presumed CFUs of Ye wt and mutant strain in feces of GF mice (upper diagram). The immune responses in GF animals are less potent as compared to C57BL/6J wild type mice while microbial CR is absent (bottom diagram). **(E)** In SPF-colonized *MyD88*^-/-^ mice we assume that the strongly limited immune reaction does not significantly affect the CR that is mediated by the endogenous microbiota. This will presumably result in a lower overall Ye cell count in the gut compared to the SPF wild type and GF mice. The immune deficiency entails an almost contingent infection outcome (right panel) resulting in either comparable numbers of the Ye wt and the mutant strain or one of the strains being more abundant at two days after infection. Please note that the infection course in the *MyD88*^-/-^ mice has to be monitored for a shorter period due to adherence to animal welfare regulations. **(F)** The presumed coincidental CFU development in feces is illustrated by overlapping, shaded areas (upper diagram). Limited immune responses are reducing CR to a low level (bottom diagram).

Consequently, Ye can colonize and replicate in the mucosal compartment if able to resist the host immune defense. As this compartment has a specific capacity, Ye cells that exceed this capacity drain into the lumen and finally end up in measurable CFUs in feces. Since only the Ye wt strain can cope adequately with the attack of phagocytes, both a YadA- and a T3SS-deficient strain will be quickly eliminated despite initial colonization as experimentally observed by us (**Fig. 1**) and others (Cornelis, 2002; Pepe et al., 1995; Ruckdeschel et al., 1996).

The situation is different in a host lacking a microbiota (**Fig. 2C and D**). In the absence of competing microbiota in GF mice, both the Ye wt and the mutant strain can expand within the lumen of the GIT (which will be termed the luminal compartment in the following) without the need to enter a mucosa-near site to colonize the gut (**Fig. 2D**). The innate immune response to Ye is presumably less intense than in SPF mice, because the immune system is not developed correctly in the absence of microbiota (Macpherson & Harris, 2004; Round & Mazmanian, 2009) (**Fig. 2D and Fig. S2**). Still, mutant strains will be eliminated more efficiently compared to Ye wt. This might lead to a slow reduction of the mutant strain at late time points after infection. Taken together, due to the lack of microbiome in GF mice, we assume that both, the Ye wt and the mutant strain, will colonize the GIT at high numbers.

In an immunodeficient host (i.e., *MyD88*^-/-^ mice), harboring a diverse microbiota (**Fig. 2E and F**), we expect a faster progression of infection (Bhinder et al., 2014; Friedrich et al., 2017; Gibson et al., 2008; Lebeis et al., 2007). Additionally, we anticipate an amelioration of the difference between Ye wt and mutant strain CFUs during the infection course, because a better colonization of Ye wt is mostly the result of its ability to survive the host immune reaction. As the immune system is only weakly active here, having the full capacity of immune evasion mechanisms is no longer a clear advantage for the Ye wt strain. Thus, we hypothesize that coinfection can result in different outcomes (wild type + mutant, only mutant, or only wild type detectable).

**In step 3**, based on our hypotheses described above, we devised the most critical interactions among Ye, the host immune system and the microbiota upon host entry and their impact on Ye population dynamics. These interactions will later be included in the model and described mathematically:

(A) Population dynamics in the mucosal compartment: after oral coinfection with a Ye wt and a mutant strain, both enter the lumen of the small intestine (SI). A portion of these Ye is then able to enter an extra-luminal location, the “mucosal compartment.” If not attached to surfaces within the SI, bacteria will be transported towards the colon due to peristalsis. Within the colon, water will be reabsorbed from the intestinal content, and all bacteria finally end up in feces. Both the retention time and the replication rate of the bacteria determine how many bacteria will be detected in feces at a distinct time point. As Ye cells presumably have a lower replication rate than the endogenous microbiota, their CFU in feces would rapidly decline compared to that of the commensals should they fail to establish a replicating population within the SI. However, experimental data show that the Ye CFU per g of content in the SI at a later time point of infection (7 dpi) is relatively high, especially in the distal part of the SI (**Fig. S4**), and we have hints that there actually does exist a niche within the GIT that can be colonized by Ye (**Fig. S5**). We hypothesize that Ye located in the mucosal compartment can resist removal by peristalsis and can even replicate. Since this compartment would have a restricted capacity only, one model assumption is that all Ye cells exceeding this capacity will re-enter and feed the luminal populations and contribute to the CFU in feces.

Bacterial interactions in the mucosal compartment: Ye expresses several virulence factors that facilitate efficient immune evasion. This capability is especially important in the mucosal compartment, where the number of immune cells and the concentration of AMPs are high. Therefore, we assume that Ye can proliferate in the mucosal compartment, which is also colonized by a small number of commensal bacteria. The growth dynamics of both Ye and commensal bacteria are determined by their initial numbers and their distinct growth rates. We assume that the endogenous microbiome in total has a higher growth rate compared to Ye because the microbiome members are rather diverse and do not necessarily compete for nutrients or suitable niches. Importantly, the combined number of all bacterial populations in the mucosal compartment is restricted by a fixed capacity. Hence, Ye and members of the microbiota compete for colonization of this compartment, and further expansion of the population is only possible if the capacity limit has not been reached yet.

(B) The influence of the immune system: host immunity involves humoral and cellular factors. For the sake of simplicity, we summarized all host defense activities in one abstract immune action that only affects the mucosal compartment, but is negligible in the luminal compartment. We hypothesize that the presence of Ye in the mucosal compartment activates the immune system. This activation increases proportionally to the number of Ye cells. As only the Ye wt strain has a full arsenal of virulence factors that allow efficient immune evasion, we assume that the Ye mutant strains and commensal bacteria are much more susceptible to killing by the immune system than the wild type.

(C) Population dynamics in the luminal compartment: most of the Ye applied orally during the initiation of infection enter the luminal compartment already populated with microbiota. We assume the same bacterial growth rates in the luminal and mucosal compartment and set a limit to the total bacterial capacity of the lumen. Moreover, this capacity is conceivably larger than that of the mucosal site. The CFU of Ye in the luminal compartment over time is - as in the mucosal compartment - determined by the initial quantity of Ye and a distinct growth rate. Additionally, bacteria that exceed the capacity of the mucosal compartment spill over into the luminal compartment and thereby contribute to the CFU in the lumen. We summarized and depicted all our considerations in **Fig. 3**.

**Figure 3.**
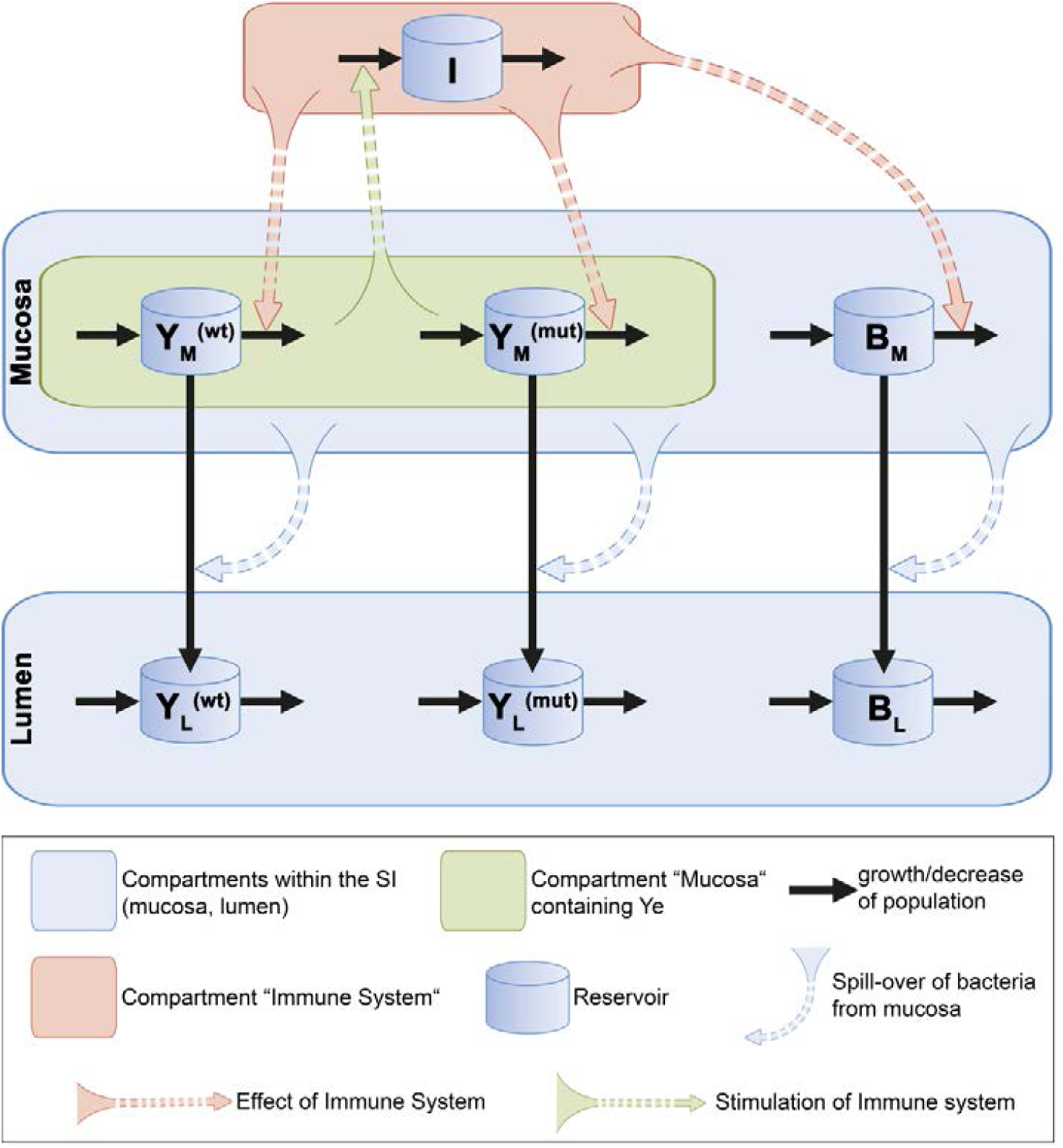
Schematic depiction of the model composition and interaction networks. The model calculates population dynamics of the Ye wt (Y_L_^(wt)^; Y_M_ ^(wt)^) and mutant strain (Y_L_ ^(mut)^, Y_M_ ^(mut)^) as well as of commensal bacteria (B_L_; B_M_) at two different sites of the small intestine (SI), the luminal site and the extra-luminal mucosal site (“mucosa”; “lumen”). Additionally, it includes an abstract immune response with a distinct immune cell population (I). Bacterial and immune cell populations are illustrated as reservoirs. Individual growth rates determine the growth of bacterial populations. Decrease of populations is caused by intestinal peristaltic movement in the lumen and by immune killing in the mucosa. Furthermore, a movement of bacteria from the mucosal compartment to the luminal compartment takes place. Upon entry of Ye wt or mutant strains to the mucosal compartment, they stimulate an immune response, which reciprocally affects all Ye and commensal populations within this compartment. Equipped with immune evasion factors, the Ye wt strain is less affected by the immune response than the Ye mutant strain, whereas both are more resistant than the commensal bacterial population (B_M_). Replicating populations that exceed the limited capacity of the mucosa drain into the lumen and thereby feed luminal populations. As a result of these bacterial population dynamics in the lumen, the model output is the calculated CFU of the bacteria ending up in feces. These curves are equivalent to experimental CFU data generated from feces of orally infected mice.

Based on the experimental data and theoretical considerations (Step 1 to 3), in **step 4**, we come up with the following mathematical model. As pointed out above, we assume that, following oral infection, a 1:1 mixture of the Ye wt and the mutant strain enters the SI. Most of the Ye remain in the lumen, but a small number enters the mucosal compartment. We assume that few commensal bacteria already populate this location. The growth dynamics of the commensal bacteria *B*_*M*_, the wild type *Y*_*M*_^(*wt*)^, and the mutant strain *Y*_*M*_ ^(*mut*)^ in the mucosal compartment are determined by their quantities and by their growth rates, described by a logistic growth with a maximum possible size. The growth rate *α*^(*B*)^ of the endogenous commensal bacteria is presumably higher than the Ye growth rates *α*^(*wt*)^ and *α*^(*mut*)^, respectively.

Moreover, the growth rates *α*^(*wt*)^ and *α*^(*mut*)^ are assumed to be equal. The capacity *C*_*M*_ limits the expansion of the bacterial population in the mucosal compartment. When bacteria counts exceed this capacity, bacteria spill over to the lumen at the following rates *σ*:

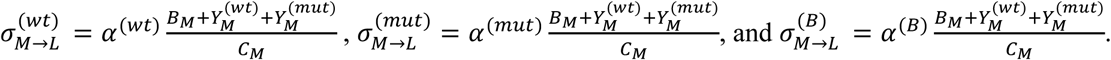

A variable that determines the infection course is the effect of *I*, the immune system. In the presence of Ye in the mucosa *Y*_*M*_, *I* is stimulated at rate *κ*, but its strength is limited to a capacity *C*_*I*_, resulting in a logistic growth 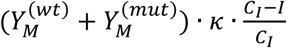.

*I* directly acts on bacteria present in the mucosa but influences only indirectly the luminal populations by affecting the spillover from the mucosal compartment into the lumen. The immune system kills *Y*_*M*_^(mut)^ more efficiently than *Y*_*M*_^(wt)^, which has a full arsenal of virulence factors that allow efficient immune evasion. However, members of the commensal microbiome *B*_*M*_ are the most susceptible to killing by *I*. This killing is modeled by using the term (*γ*· *I* · *B*_*M*_). We use the adjustment factors 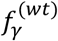 and 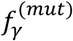 to account for the different susceptibilities of *Y*_*M*_^(wt)^ and *Y*_*M*_^(mut)^ towards killing by *I* and the even higher susceptibility of *B*_*M*_. The following differential equations describe the resulting dynamics of bacterial populations and immunity strength at the mucosal site:

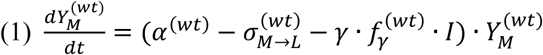

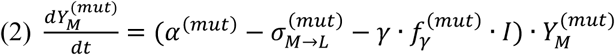

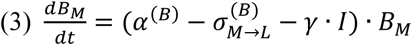

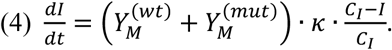

Most of the Ye from the oral infection enter the lumen of the SI. Additionally, luminal populations are fed by bacterial spill over from the mucosal compartment. The lumen is already populated with commensal bacteria. For the sake of simplicity, we use the same bacterial growth rates *α*^*(B)*^, *α*^*(wt)*^, and *α*^*(mut)*^ in the lumen as at the mucosal site. As we limit the total bacterial capacity of the lumen to a large number *C*_*L*_, we obtain the following logistic growth for the luminal compartment:

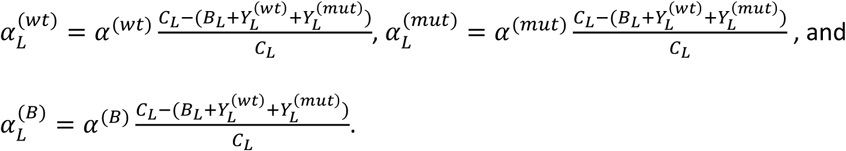

Bacteria in the lumen move along the intestinal tract and are finally excreted at a removal rate *β*. Combining all this, the following set of equations gives the dynamics of the bacterial populations in the lumen:

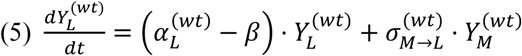

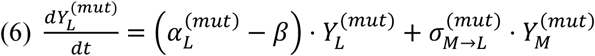

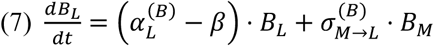

One of the most challenging steps in modeling is the estimation of unknown parameter values in an ordinary differential equation system from experimental data **(Step 5)**. In order to solve the system, we, therefore, aimed to reduce the number of parameters with unknown values. This was achieved either through experimental approaches, if possible, by estimating biologically meaningful ranges for unknown parameters (based on literature and own data), or, at least, by defining the relations between distinct parameters (higher/lower/same as). To this end, we experimentally determined the gut passage time of C57BL/6J wild type SPF (termed SPF from now on), C57BL/6J wild type GF (termed GF from now on), and *MyD88*^*-/-*^ SPF (termed *MyD88*^*-/-*^ from now on) animals and found that in the GF animals the gut passage time is much longer than in SPF and *MyD88*^*-/-*^ animals (**Fig. S6**). We also determined immunological parameters of SPF, GF, and *MyD88*^*-/-*^ animals, thus supporting our assumptions in regard to the relative strength of the immune response in the three distinct systems (**Fig. S2**).

To find reasonable values for parameters that either cannot at all be determined experimentally or only with non-justifiable cost and effort, we started a computational parameter optimization to yield predictions in best agreement with experimental data. Therefore, we used built-in optimization methods of MATLAB (see Materials and Methods). Detailed information for all parameters (definition, source of parameter values, function, and relation to other parameters) is given in **Table 1**. Of note, the model implementation and the optimization process were at first based on the dataset generated from the coinfection of SPF wild type mice with the Ye wt and the YadA0 mutant.

**Table 1.**
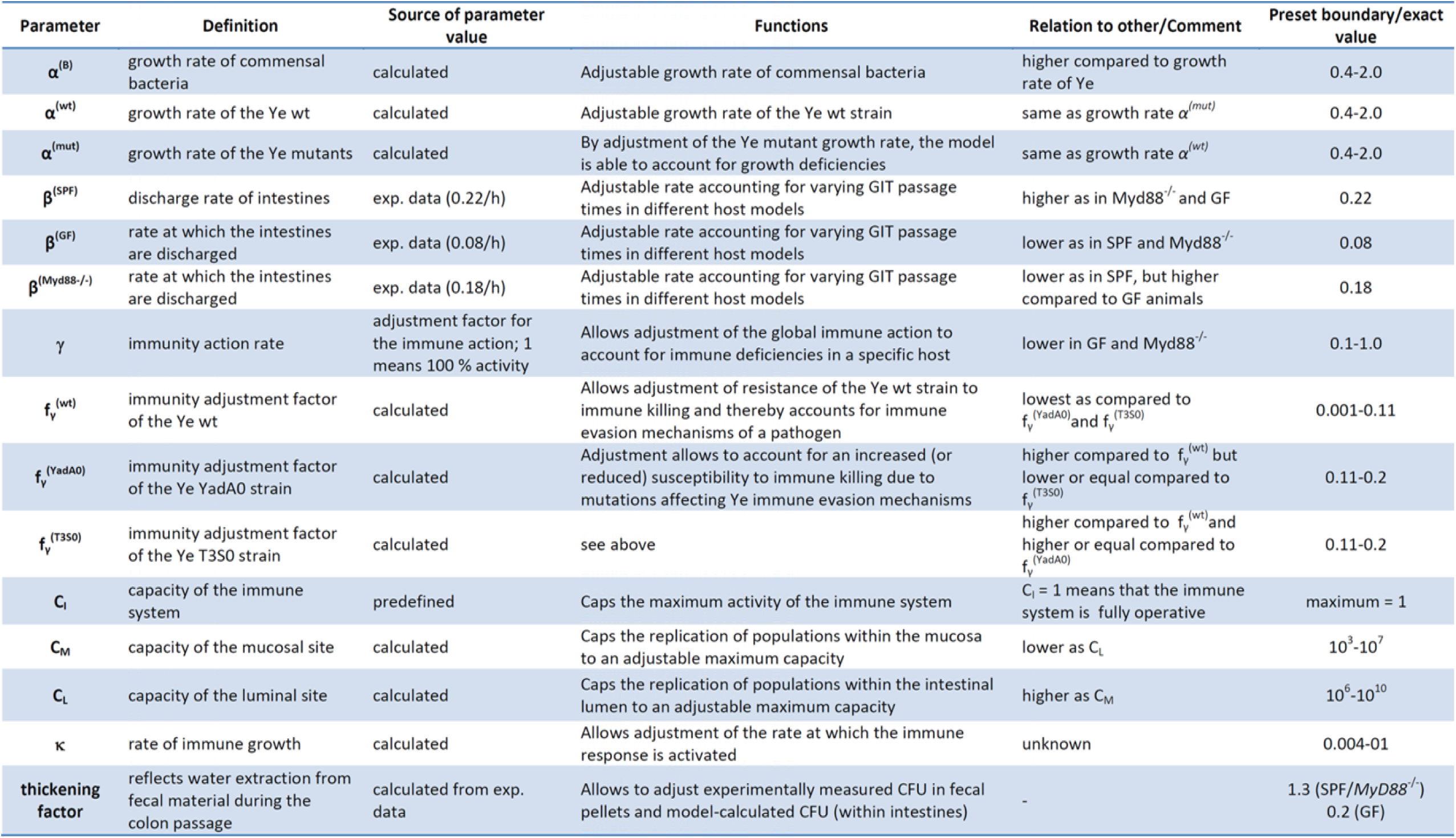
Overview about model parameters, source of values, function, relation to other parameters and preset boundary or the exact value used for parameter calculation

When evaluating the model (**Step 6**), we found that the model output was in good agreement with the Ye population dynamics that we determined experimentally in SPF mice co-infected with Ye wt and Ye YadA0 (**Fig. 4A)**. Independent estimation of parameters based on a second experimental dataset that was obtained by co-infection of SPF mice with Ye wt and Ye T3S0 delivered slightly different, but comparable absolute parameter values compared to Ye wt : Ye YadA0 coinfection. Hence, we did observe concordance of the model output with the experimental data (**Fig. 4B**). Strikingly, the model even reflects a difference between the dynamics of CFU development of the Ye YadA0 and the Ye T3S0 strain.

**Figure 4.**
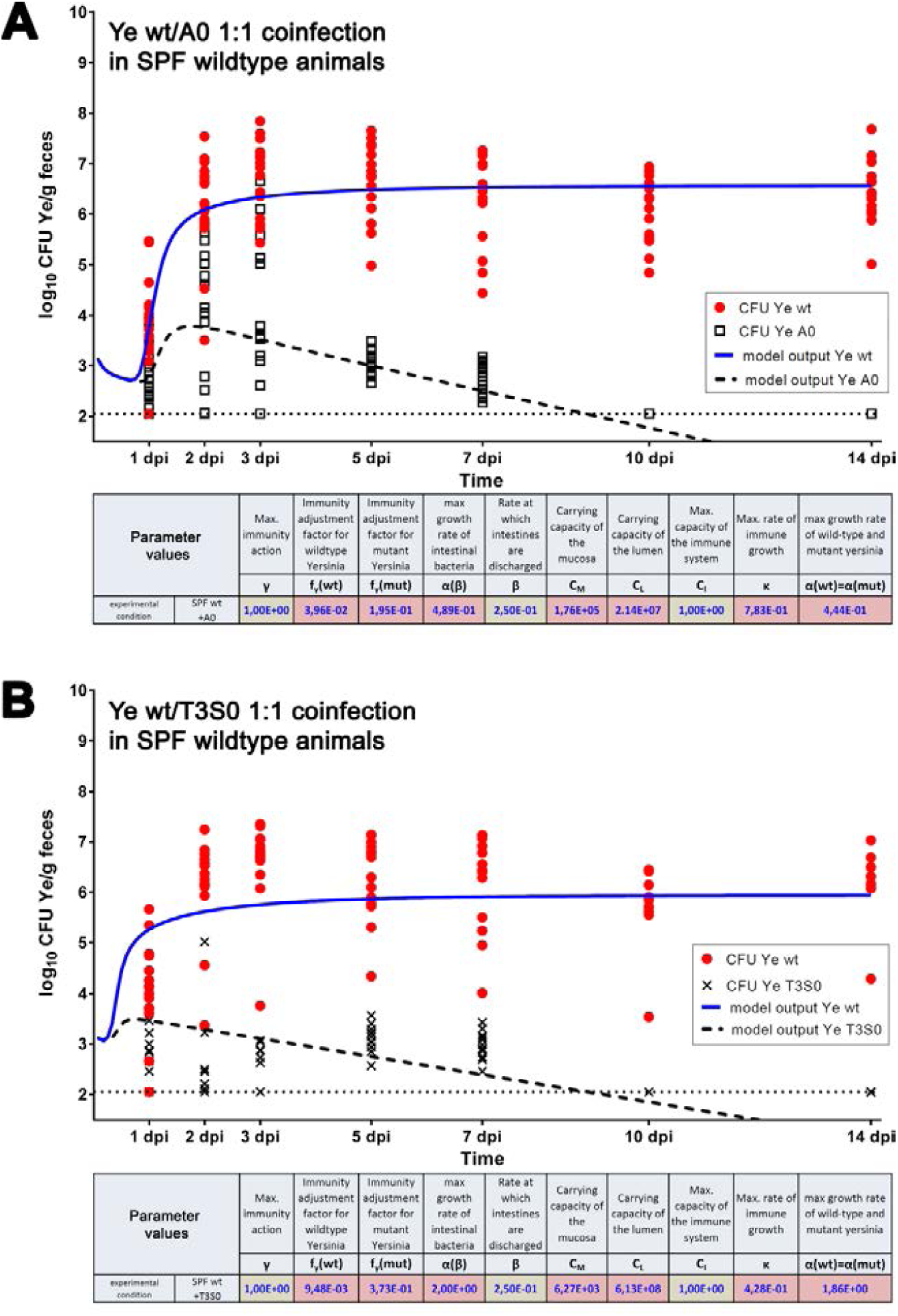
Overlay of model output and experimentally determined CFU values during Ye coinfection of SPF wild type mice. For the model prediction, the listed parameter values were used. **(A)** Model output for CFU of Ye wt and Ye YadA0 shown as an overlay with experimental data. CFU values of individual animals at indicated time points are shown for Ye wt and Ye YadA0. The dotted line indicates the limit of detection of our experimental system. **(B)** Model output for CFU of Ye wt and Ye T3S0 as an overlay with experimentally determined CFU values from the Ye wt : Ye T3S0 coinfection of SPF wild type mice. Calculated parameter values (red background) and fixed parameter values (green background) are shown in the tables.

Moreover, our finding that Ye T3S0 is more susceptible to killing compared to Ye YadA0 is also corroborated by the model. Looking at the relative values compared to Ye wt 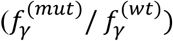, the Ye YadA0 strain is ∼5 times and the Ye T3S0 ∼ 40 times more susceptible to killing by the immune system. The calculated parameter values obtained for these experimental datasets are depicted as insets in **Fig. 4**.To better comprehend how changes in the relations of 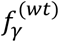 and 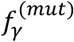 impact on CFU development we additionally created **Fig. S7**. Taken together, the predicted population development in the SPF wild type host for both coinfection settings served as a proof of the appropriateness of the model as it proved to be in line with the experimentally observed infection course.

### Challenging the model: Lack of microbiota

In order to challenge our model, different basic parameter settings for microbiota-derived CR and host immune competence were adapted, and the resulting model predictions were analyzed by comparing them to experimental coinfection data. To decipher the effect of the absence of the microbiota on CFU development, we defined the number of *B*_*M*_ and *B*_*L*_ (i.e., number of bacteria in mucosal (M) and luminal compartment (L)) to be 0. Moreover, we considered that the fecal pellets have a higher water content in GF mice, as experimentally determined (**Table S8**). The higher water content was reflected by using a different thickening factor. Furthermore, we took into account the lower discharge rate in GF mice (12 h mean residence time instead of 4.5 h in SPF animals) which we had also determined experimentally (**Fig. S6**). Experimental coinfection of GF mice with Ye wt + Ye YadA0 or Ye wt + Ye T3S0, respectively, revealed that both the Ye wt and the mutant strains reached remarkably higher cell counts in feces as compared to CFU levels in SPF colonized mice. The T3S0 strain exhibited a slight attenuation resulting in apparently lower CFUs, particularly from 7 dpi on, whereas Ye wt and Ye YadA0 counts remained constant at a high level over the entire observation period of 14 days (**Fig. 5**). Our data thus indicate that in the absence of a commensal microbiome, both YadA and the T3SS seem to be dispensable for effective colonization of the GIT.

**Figure 5.**
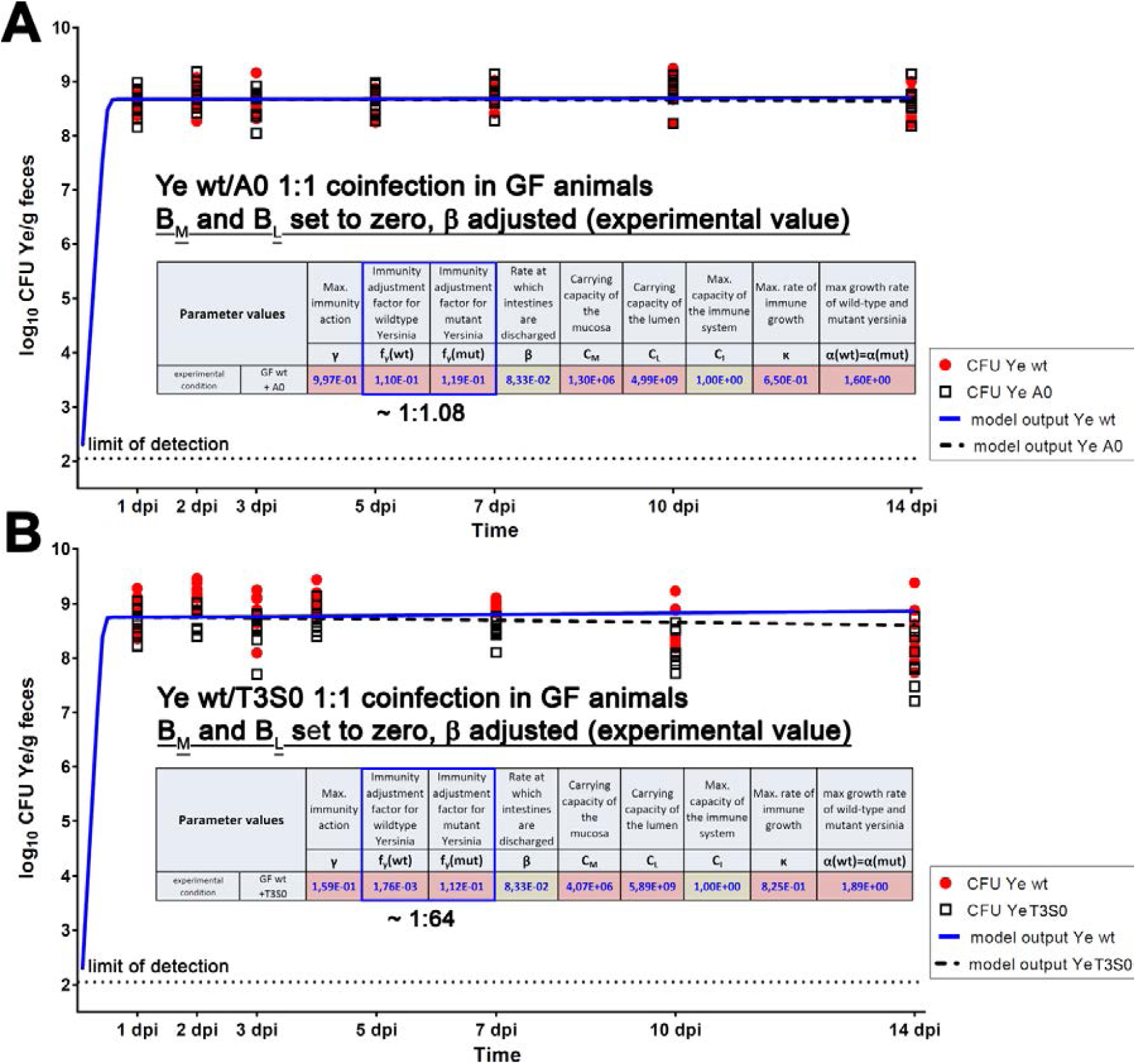
Challenging the model part I: Absence of microbiota. **(A)** Overlay of model output for CFU of Ye wt and Ye YadA0 or **(B)** Ye wt and Ye T3S0 and experimentally determined CFU levels from coinfections of GF mice. All parameters were estimated based on respective experimental data (parameter values are listed in the inset table).

Next, we ran the model for the Ye wt : Ye YadA0 coinfection setting by keeping defined boundaries only for some parameters that were justified from a biological point of view (**Table 1**), and values we had determined experimentally. The model output was in good agreement with the experimentally determined course of CFU development (**Fig. 5A**). We also found that the parameter values that the most differed from what we had previously obtained for the SPF wild type model were higher capacities C_M_ and C_L_ for the mucosal and the luminal compartment, respectively. This makes sense, as GF animals have massively enlarged intestines. Interestingly, 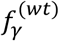 and 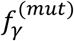 were estimated to be very similar (0.110 for Ye wt and 0.119 for Ye YadA0). This corroborates our interpretation of the infection course in GF mice. Here, the Ye YadA0 strain does not have any disadvantage compared to the Ye wt strain and can expand within the gut to the same extent. Thus, also in the model, YadA seems to be dispensable for effective colonization in the absence of a microbiota.

When we estimated *γ* in this setting, we obtained an optimized value of ∼ 0.997 (**Fig. 5A**), which is very similar compared to SPF. This finding was surprising as we had expected lower activity of the immune system in the GF setting according to literature and our own data (see also **Fig. S2**). However, our model predicts that the overall influence of *γ* on the expansion of Ye is only subtle (**Fig. S9**). This can be explained by the absence of the endogenous microbiota that competes with Ye for filling the capacity of the small intestine in the SPF animals. We also modeled the GF Ye wt : Ye T3S0 coinfection and obtained very similar results compared to the Ye wt : Ye YadA0 coinfection (**Fig. 5B**). The most apparent difference was that the predicted CFU for the T3S0 mutant strain slightly dropped towards the end of our observation period, which is in line with our experimental data. Again, this difference in the behavior of Ye YadA0 and Ye T3S0 can be explained with their different susceptibility to killing by the host immune system. As in the absence of a microbiome both strains can expand very quickly, the effect of the increased susceptibility of Ye T3S0 to killing is not as high as in the SPF model system. Taken together, our model can compute Ye population dynamics also under GF conditions, and the results correlate with our experimental data.

### Challenging the model: The immunocompromised host

As a second evaluation of our model, we aimed to mimic an immunocompromised host. We made use of *MyD88*^-/-^ C57BL/6J mice that were colonized with a SPF microbiota as a model to decipher the role of a restricted immune response in Ye population dynamics. We assumed a more rapid and frequent invasion due to the reduction of the immune response, as depicted in **Fig. 2E** and **2F**. As in the SPF wild type model, in the *MyD88*^*-/-*^ animals, Ye encounters the mucosal compartment occupied by commensals. Because of the MyD88 deficiency, a much weaker immune response is induced. This primarily has two consequences: (i) The microbiota is less disturbed and reduced. Therefore, Ye is less successful in establishing a population in the mucosal compartment, and the Ye counts will be lower. As the mucosal compartment feeds the luminal Ye population by spill over, we will observe a lower Ye CFU in the GIT compared to C57BL/6J wild type animals. (ii) Due to the weak immune response of the *MyD88*^*-/-*^ animals, we assume that the disadvantage of the mutant strains in competition with Ye wt is much less pronounced.

Finally, we co-infected SPF colonized *MyD88*^*-/-*^ mice, as described before. To compare the experimental results and modeling data, we created an overlay of the model output and the experimental data (**Fig. 6**).

**Figure 6.**
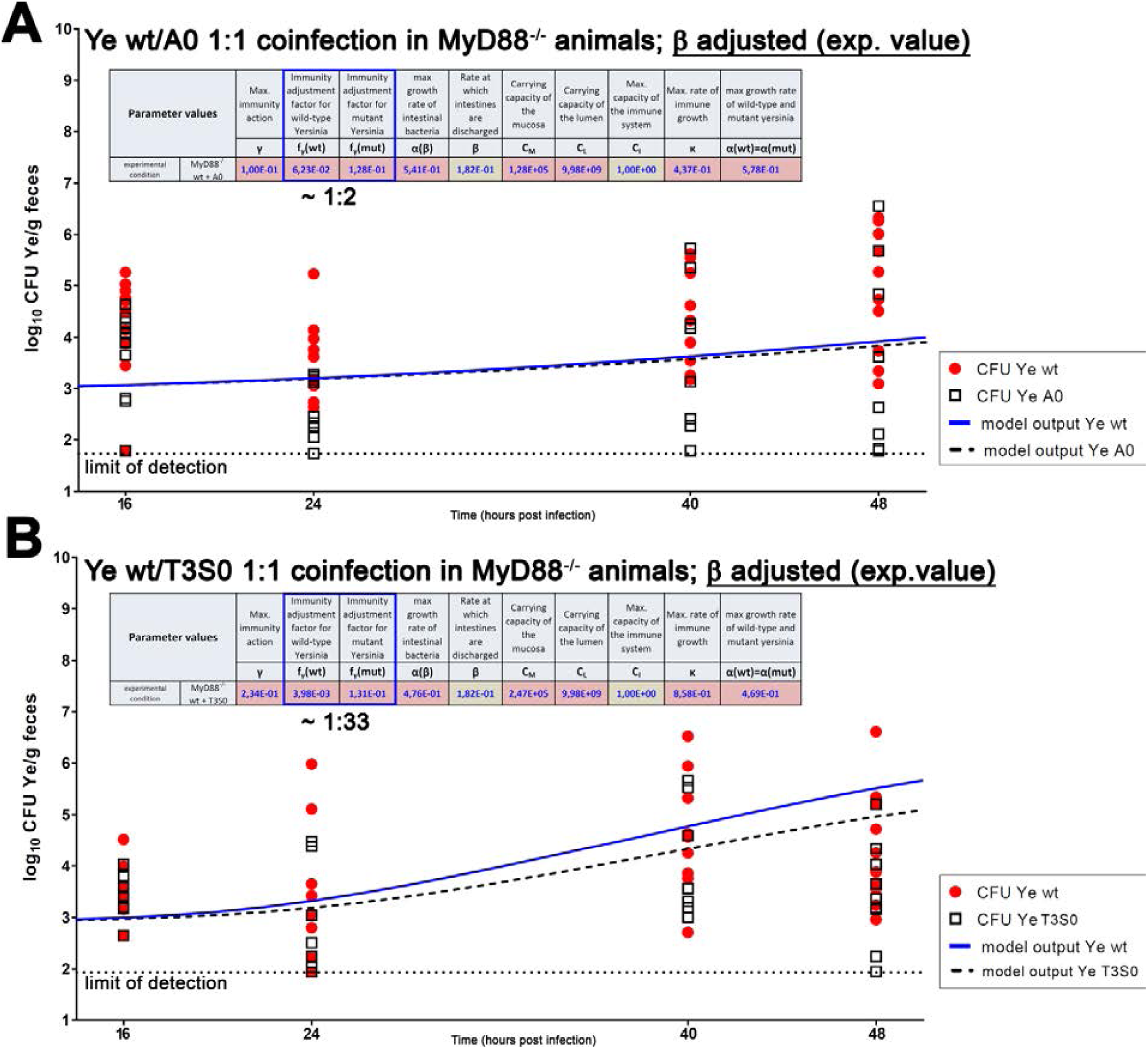
Challenging the model part II: Impaired immune response (*MyD88*^*-/-*^) **(A)** Overlay of model output and experimentally determined CFU levels from coinfections of SPF *MyD88*^*-/-*^ mice with Ye wt and Ye YadA0 and **(B)** Ye wt and Ye T3S0. All parameters were estimated based on the respective experimental data (parameter values are listed in the inset table).

Due to the high frequency of fatal infections that has been observed with *Salmonella* Typhimurium and *Citrobacter rodentium* (Bhinder et al., 2014; Gibson et al., 2008; Hapfelmeier et al., 2005), infections with Ye were conducted for two days only. To get a better temporal resolution within this shorter observation period, the Ye counts in feces were determined at two additional time points (i.e., after 16 and 40 h). Within 48 hours post-infection, the CFU of Ye wt showed a slight increase compared to earlier time points but as expected never reached as high counts as we had observed in SPF wild type mice. The mean CFU of Ye YadA0 was marginally lower compared to that of Ye wt (**Fig. 6A**), whereas the difference in CFU of Ye wt compared to Ye T3S0 was more pronounced, but also subtle (**Fig. 6B**). In some of the *MyD88*^-/-^ mice, the YadA0 and, to a lesser extent, the T3S0 strain reached a comparable, or even a higher CFU than the one of Ye wt strain at 48 hpi.

In summary, we found that (1) our model is appropriate to predict the infection course in immunocompromised animals, (2) that a proper immune response outreaches the importance of the presence of the microbiome in preventing colonization and infection with Ye, and (3) that both YadA and the T3SS seem to play only a minor role in the colonization of the GIT.

## Discussion

The complex interplay of a specific pathogen with host factors, as well as the integrity and composition of the endogenous microbiome, determines the outcome of a gastrointestinal infection. Herein, we developed a model to simulate the dynamics of bacterial populations in enteropathogenic infection and to predict the infection course.

Our main findings are that the model can predict the infection course in different host settings (immune-competent host with a diverse microbiota, no microbiota, immunocompromised). However, each setting involves its own distinct parameter set. To predict the infection course reliably, it was not enough to alter individual parameters to adopt a change implied by a specific condition (e.g., no microbiota present). Only if parameter values were optimized based on the respective experimental dataset, the predictions were in good agreement with our experimental observations. Presumably, the differences in structural and functional details (e.g., GIT morphology and physiology, gut passage time), even in our basic experimental setting (comparing SPF and GF animals) entail that the parameter values are not merely exchangeable between systems. Within consistent host condition and pathogen phenotype, however, the infection course can be predicted mathematically.

We conclude from our study that an excellent understanding of the causative agent of GIT infection is needed: How does the pathogen interact with the host? Does it produce specific virulence factors? How do these factors contribute to population dynamics (e.g., by mediating immune evasion)? Does the pathogen have specific requirements for growth (e.g., oxygen, nutrients)? These and many more questions need to be known or clarified experimentally in the best case. Consequently, our current model can in principle be used to predict the infection course of other pathogens, but needs to be adapted by concerning their specific peculiarities with regards to the above mentioned characteristics. Such adaptations might be easily done with pathogens that have a lifestyle comparable to that of Ye, but will require profound changes of the model setup for other pathogens.

To create a model that delivers rational predictions, we also need a good understanding of the infected host: is its microbiota able to mediate full colonization resistance? Was the microbiota already disturbed by medication? Is the immune system fully operable? Is the GIT physiology disturbed (leading to, e.g., prolonged or impeded gut passage)? The more detailed our understanding of the pathogen and the host, the better the model can reflect biology.

In recent years, several mathematical models were developed to mirror bacterial gastrointestinal infections (Grant et al., 2008; Jones & Carlson, 2018; Kaiser et al., 2014; Kaiser et al., 2013; Leber et al., 2017; Verma et al., 2019), viral infections at epithelial sites (Miao et al., 2010), and inflammatory disorders such as IBD (Balbas-Martinez et al., 2018; Wendelsdorf et al., 2010). Our model emphasizes the trilateral relationship of the enteropathogen, the host and the microbiota. By enabling the modulation of pathogen and host specific properties, it significantly contributed to extend our knowledge about their role for the course of infection.

There are several aspects that should be included in more refined versions of our model: (I) A better reflection of the growth dynamics of both the pathogen and the microbiota within the GIT. Different approaches could resolve this issue (Grant et al., 2008; Jones & Carlson, 2018; Myhrvold et al., 2015; Simůnek et al., 2012; Stein et al., 2013). Other groups have addressed bacterial colonization dynamics in the intestines and translocation events after *Salmonella* Typhimurium infection. They parametrized their mathematical models with data from tagged isogenic *S*. Typhimurium strains (Grant et al., 2008; Kaiser et al., 2013; Moor et al., 2017). The adoption of this methodology to the Ye infection model could provide more detailed insights into the population dynamics at specific sites, such as the mucosal compartment.

Another desirable amendment of the model would be (II) the implementation of a more sophisticated immune system to increase the flexibility of the system. Several studies detailed the complex network of the host immune response that is activated by a given pathogen (Balbas-Martinez et al., 2018; Leber et al., 2017; Miao et al., 2010; Verma et al., 2019; Wendelsdorf et al., 2010). The results of these studies could be used for implementation, of course making the model system much more complicated.

In order to assess how well the model can be adapted to other pathogens, it would be desirable to investigate its performance in predicting the infection course of other clinically relevant species, such as enteropathogenic *E. coli*. However, this was beyond the scope of this study and needs to be investigated in the future, together with experts in the field having the required knowledge about the respective infection biology and the access to suitable animal infection models.

Finally, it would be great to include (IV) the possibility to reflect external perturbations such as the treatment of the host with specific antibiotics. This could possibly be achieved by integrating data about the resistance phenotype of the pathogen and the impact of changes in microbiota composition and colonization resistance. The dynamics of the intestinal microbiota composition have previously been addressed in modeling approaches, especially in the context of *Clostridium difficile* infection. Time-dependent metagenomics data were used to analyze the influence of antibiotic perturbations on microbiota and pathogen overgrowth *in silico* in an approach that combined a Lotka-Volterra model of population dynamics and regression (Buffie et al., 2015; Stein et al., 2013). A recent extension of this model incorporated antibiotic resistance mutations and sporulation as further virulence attributes of *C. difficile* (Jones & Carlson, 2018). An adaption of this specific model could, in the future, lead to more elaborate model predictions in terms of microbiota perturbations due to antibiotics.

In sum, we think that computational modelling of infection has a great potential, but also many caveats. It is tempting to speculate whether at some point computational modeling could be used to predict the infection course of patients at risk and if such predictions could really be used to improve patient treatment and outcome. Main caveats are the huge complexity of the system “patient”, and also the plasticity of the causative pathogens. Of course, we are aware that our results using an animal model are merely transferrable to the human hosts. Still, we hope that with our study, demonstrating the feasibility and usefulness of infection modelling we have contributed a small piece to make this true in the far future.

## Materials & Methods

### Bacterial strains and growth conditions

Ye wt and mutant strains used in this study are listed in Supplementary **Table S10**. All strains were cultured overnight at 27°C in Luria Bertani broth (LB). As selective antibiotics nalidixic acid (10 µg/ml), kanamycin (50 µg/ml), spectinomycin (100 µg/ml) and chloramphenicol (25 µg/ml) (all Sigma-Aldrich) were supplemented in combinations according to the indicated resistances (**Table S10**). For the preparation of bacterial suspensions for oral infection, overnight cultures were diluted and subcultured for 3 h at 27°C. Bacteria were then washed once with Dulbecco’s phosphate-buffered saline (DPBS, Gibco, Thermofisher) and the OD_600_ was determined to prepare the desired inoculum.

### Generation of Ye strains containing different antibiotic selection markers

A Chloramphenicol resistance cassette derived from pASK IBA4C (IBA Lifesciences) was chromosomally introduced into the YenI locus of the Ye WAC strain to discriminate between the Ye wt and the Ye YadA0 or the T3SS deficient strain (T3S0). The YenI gene encodes for a Ye specific restriction-modification system the interruption of which allows higher efficiency of genetic manipulations (Antonenko et al., 2003; Miyahara et al., 1988). The resistance cassette was inserted by homologous recombination using the suicide plasmid pSB890Y as described previously (Weirich et al., 2017), and insertion was verified by PCR, antibiotic resistance testing and sequencing. Finally, the respective virulence plasmids were re-transformed into Ye WAC Cm^R^. All plasmids and primers used for the insertion of selection markers are listed in the Supplementary Tables S1 and S2.

### Animal handling

Ethics statement: all animal infection experiments were approved by the regional authority of the state Baden-Württemberg in Tübingen (permission number H2/15). Female C57BL/6J OlaHsd mice were purchased from Envigo (Horst, NL). MyD88-deficient mice (*MyD88*^-/-^) with C57BL/6J genetic background were obtained from a local breeding colony (breeding pairs were purchased from Jackson Laboratories). Animals were housed in the animal facility of the University Hospital Tübingen under specific-pathogen-free (SPF) conditions. Germ-free (GF) animals were bred in the germ-free core facility of the University Hospital Tübingen or provided by the Institute for Laboratory Animal Science (Hannover Medical School, Germany). All animals were housed in individually ventilated cages in groups of 5 animals and were supplied with autoclaved food and drinking water *ad libitum*. Infection experiments were performed with female mice at 6-10 weeks of age.

### Oral mouse infection

Prior to the intragastric administration of bacteria, mice were deprived of food and water for 3-4 hours. For oral coinfection experiments, animals were infected with a 1:1 mixture of each 2.5·10^8^ CFU of Ye wt and Ye YadA0 or Ye T3S0, respectively. Upon oral coinfection, SPF wild type and GF mice were sacrificed at time points indicated within the figures describing the results of individual experiments. *MyD88*^-/-^ mice were infected for two days only because of the expected rapid systemic spread in these immunocompromised animals. Oral infections for subsequent RNA analyses from small intestinal mucosal scrapings were performed for two days.

### Determination of bacterial load from feces

Fresh fecal pellets were collected after manual stimulation of individual mice, weighed, and resuspended in 500 µl sterile DPBS. Pellets were homogenized, serially diluted with DPBS, plated on selective agar plates, and incubated at 27°C for 48 h. Afterwards colonies were counted, and the CFUs per gram of feces was calculated.

### Calculation of competitive indices in mixed infections

Competitive indices (CI) from fecal and tissue samples were calculated as the CFU output of the Ye mutant/Ye wild type strain divided by the input (initial oral inoculum) of these strains (CFU Ye mutant strain input/CFU Ye wild type strain input) (Dyszel et al., 2010). The output was determined in the individual experiments as described above. The initial oral inocula (= the input) were verified by serial dilution and subsequent plating on LB with appropriate antibiotics. A CI with a logarithmic value of zero indicates identical fitness of the wild type and the mutant strain, while a negative CI indicates that the mutant strain is impaired in colonization (Dyszel et al., 2010).

### Isolation of RNA from gut mucosal scrapings

For isolation of total RNA from gut mucosal scrapings, five mice per group harboring either SPF microbiome or GF and the genetic backgrounds indicated earlier were infected with a 1:1 mixture of each 2.5·10^8^ CFU of Ye wt, and Ye T3S0. As controls, five mice of each colonization state and genetic background were orally administered with 100 µl PBS instead of bacterial suspensions. Two days after infection the mice were sacrificed and the distal 10 cm of the small intestine was dissected and shortly incubated in RNA later (Qiagen). Then the tissue was flushed with ice-cold DPBS to remove the fecal content and opened longitudinally on ice using scissors. After the removal of residual feces by flushing again with ice cold DPBS, the mucosa was scraped off with the blunt side of a scalpel and incubated overnight in RNA later at 4°C. RNA later was removed, and scrapings were homogenized in TRI-Reagent (Zymo Research) by rinsing them successively through syringe needles with decreasing diameters. The remaining cell debris was removed by centrifugation, and the supernatants were finally used for the RNA purification using the DirectZol RNA Miniprep Plus Kit (Zymo Research) according to the manufacturer’s protocol. This protocol included a step for the removal of contaminating genomic DNA. The resulting RNA was quantified using a Nanodrop photometer (Thermo Fisher), and the integrity was confirmed by agarose gel electrophoresis.

### Quantification of immune parameters by quantitative real-time PCR (qRT-PCR) (Figure S2)

Relative mRNA levels of target genes were determined using qRT-PCR. After an additional treatment for removal of genomic DNA included in the QuantiTect reverse transcription kit (Qiagen), mRNA was reverse transcribed according to the manufacturer’s protocol using 1 µg of RNA as input for a 20 µl reaction. For subsequent qRT-PCR, the TaqMan gene expression master mix (Applied Biosystems; all assays are listed in suppl. Table S2) was used with thermal cycling conditions according to the manufacturer’s protocol. cDNA input was 5 µl for a 20 µl PCR sample. Absolute quantifications were performed on a LightCycler 480 instrument (Roche) using the LightCycler 480 Software 1.5. Relative gene expression levels of target genes to the reference gene *beta-glucuronidase* (accession number AI747421) (Wang et al., 2010) were determined to apply kinetic PCR efficiency correction, according to the method of Pfaffl (Pfaffl, 2001) and normalized to the expression levels of uninfected SPF-colonized mice.

### 16S rRNA sequencing from SI luminal samples (Figure S3)

For analysis of microbiota composition within the SI of mice and to assess changes in microbiota composition upon infection with Ye, mice were initially co-housed for ten days. After oral infection with Ye as described before, or after gavage of the same volume of DPBS, mice were sacrificed at the indicated time points. The entire GIT was dissected, and the SI was removed. Intestinal contents were isolated by gently squeezing them into tubes using sterile forceps. After that, the samples were immediately snap-frozen and stored at -80°C until DNA isolation. DNA was extracted as described in the international human microbiome project standard (IHMS) protocol H (http://www.microbiome-standards.org/fileadmin/SOPs/IHMS_SOP_07_V2.pdf) (Godon et al., 1997; *IHMS Protocols*). Library preparation and 16S rRNA amplicon sequencing were performed by the CeMet Company (Tübingen) using variable regions v3-v4. Paired-end sequencing was performed on the Illumina MiSeq platform (MiSeq Reagent Kit v3) with 600 cycles. Raw read quality control was done using the FastQC tool (http://www.bioinformatics.babraham.ac.uk/projects/fastqc/) (“Babraham Bioinformatics - FastQC A Quality Control tool for High Throughput Sequence Data,”). To this end, reads were merged and quality filtering was performed using USEARCH (Edgar, 2010). Taxonomy data annotation of sequences was done by comparison against the National Center for Biotechnology Information (NCBI) bacterial 16S rRNA database using MALT (Herbig et al.). Abundance tables at the taxonomic rank of interest were generated using MEGAN6 (Huson et al., 2016) and further analyzed using the software R (http://www.R-project.org)(*R: The R Project for Statistical Computing*) (“R: The R Project for Statistical Computing,”). Before statistical analysis, all samples were normalized to 14947 reads using the tool rrarefy which is part of the vegan package (Dixon, 2003). The vegan package diversity function was used to calculate Shannon diversity. An unpaired Wilcoxon sum rank test determined significant differences between groups. Vegsdist and prcomp (also part of the vegan package) were used to perform principal component analysis (PCA) on Bray-Curtis dissimilarities. For the generation of graphical output, ggplot2 (Gómez-Rubio, 2017) was employed. 16S rRNA sequencing data will be published on the European Nucleotide Archive with the study accession number PRJEB37566.

### Determination of the distribution of Ye along the mouse GIT (Figure S4)

To determine the ratio of Ye and cultivable commensal bacteria in the different compartments of the GIT, three mice were orally infected with 5·108 CFU of the Ye wt strain. Seven days after infection, mice were sacrificed, and the gut was dissected. A piece of 1 cm length directly adjacent to the stomach was removed, and the residual small intestine was split into three pieces of equal length: a proximal part (SI 1), a middle part (SI 2), and a distal part (SI 3). Additionally, the cecum and the colon were dissected. The contents of the three small intestinal pieces, the cecum, and the colon, were isolated by gently squeezing them into tubes using sterile forceps. For each compartment, the CFU per gram intestinal content was determined as described above for feces, using selective agar to determine Ye CFUs and brain heart infusion agar (BHI; incubated in anaerobic pots) for determination of the approximate number of cultivable commensal bacteria.

### Systemic administration of gentamicin for cleansing of a potential niche colonized by Ye (Figure S5)

In order to find out about the existence of extra-luminal Ye that drain into the lumen of the SI, we tested if the systemic administration of an antibiotic that can kill Ye but is not able to enter the lumen of the GIT might reduce the Ye burden in feces. To this end, 14 mice were coinfected with Ye wt and Ye YadA0 for two days. At this time point, we assumed the successful colonization of a niche and high bacterial burden in the feces. Mice were then split into two groups, of which one was administered intraperitoneally 40 mg/kg gentamicin (Ratiopharm) in 200 µl 0.9 % sterile NaCl (Braun) and the other group sterile saline only. Ye CFUs were determined from feces of mice before gentamicin/saline administration (i.e., on 2 dpi) and one day after treatment (i.e., on 3 dpi) as described above. On 3 dpi mice were sacrificed, and Ye CFUs were additionally determined from Peyer’s patches.

### Determination of GIT passage time (Figure S6)

SPF C57BL/6 wild type or *MyD88*^*-/-*^ mice, as well as GF wild type mice (2 mice/group), were orally challenged with 100 µl DPBS containing 1·10^9^ fluorescent polystyrene beads (1 µm) (Thermo Fisher) plus 5·10^8^ CFU Ye wt in order to simulate infection conditions. After the gavage, fecal pellets were collected hourly over 24 hours, weighed, snap-frozen, and stored at -20°C until analysis. Next, samples were homogenized in 1 ml PBS and debris was removed by a centrifugation step of 20 min with 50 × g (van der Waaij et al., 1994). To determine the number of fluorescent events per gram of feces, the resulting supernatant was spiked with a defined number of compensation beads (BD biosciences) in order to be able to determine the number of fluorescent beads in a defined volume by flow cytometry. The cumulated bead-hours were then calculated by multiplying the number of beads detected by the time spent in the gut until excretion. The mean residence time per bead was finally calculated by dividing the number of summarized events/g feces by the total bead-hours.

### Determination of water content of SI content and fecal pellets (Table S8)

Three mice each, with either SPF microbiota or GF, were used for this experiment. Before dissection of the GI tract to determine the water content, 2-5 fecal pellets were collected. Then mice were sacrificed, and the entire GI tract was removed. Afterward, the stomach was discarded, and the small intestine was cut into two pieces of comparable length. Then the cecum and colon were dissected. All pieces and the fecal pellets were placed into individual, weighed Petri dishes. After that the wet weight of all samples was determined. The SI pieces, the cecum and colon were then cut open, and the content was scratched off and transferred into the Petri dish. The remaining emptied tissue was removed and weighed again, and the wet weight of the contents was determined. After that, the Petri dishes were placed without lids into an incubator, and the material was dried overnight at 65°C. Then all samples were weighed again to determine the dry weight. Finally, the total water content was calculated by subtracting the dry weight from the wet weight.

### Calculation of the thickening factor for SPF and GF mice

Our model predicts the dynamics of the number of *Yersinia* (i.e., CFU) within the SI, whereas our experimental observations are based on colony counts derived from the plating of fecal pellets (log_10_ CFU per g of feces). To align model output to experimental data, we determined the mean percentage of water in different sections of the gastrointestinal tract of SPF or GF mice and considered that the small intestinal content is massively concentrated to be excreted as a solid fecal pellet. Based on these data, we calculated a “thickening factor”. The content of the SI of SPF mice has a rather different percentage of water (77 %) compared to that of fecal pellets (29 %; **Table S8**). Therefore, the model predictions were multiplied with a correction factor in order to relate model output to laboratory observations. This factor is obtained by dividing the product of 1 g of fecal pellets and its content of solid matter (100 % - 29 %) by the product of the volume of SI content of SPF mice (which is about 2.3 g) and its content of solid matter (100 % - 77 %), i.e., the factor is (1 g · 71 %) / (2.3 g · 23 %) ≈ 1.3. Thus, our model output needs to be multiplied by 1.3 before it can be compared with experimentally determined CFU levels. GF mice differ in several aspects of SPF mice. They have a massively enlarged intestine (we measured the volume of intestinal contents to be about 10 g). The average water content of the fecal pellets is 49 % in these mice. Using the same calculation as above, we obtain a multiplication factor of (1 g · 51 %) / (10 g · 23 %) ≈ 0.2 for GF mice.

### Alignment of model simulation and lab observation time

We determined the passage of the GIT to take on average 4 h in SPF wild type mice, 5.5 h in *MyD88*^*-/-*^ mice and 12 h in GF mice (**Fig. S6**). Assuming 1 h passage time in the stomach and 1 h in the colon, this leaves a sojourn time of 2 h (3.5 h in *MyD88*^*-/-*^ mice and 10 h for GF) in the SI in which Ye are assumed to multiply. Our model only describes what is happening in the SI, starting when Ye leave the stomach (this corresponds to 1 hpi). Then an additional hour is needed for the colon passage until the CFUs can be counted. Thus, the observation in the laboratory at, e.g., 24 h after oral infection must be compared with the model results after 22 h of model simulation. This time shift of 2 h is taken into consideration whenever modeling results and experimental data are compared.

### Parameter optimization

We derived a 7-dimensional ordinary differential-equation system describing Ye population dynamics with seven and eight unknown parameter values (7 in SPF and 8 in both GF and *MyD88*^*-/-*^). These values were estimated by solving an optimization problem using the maximum likelihood method. The objective function was to minimize the Euclidean distance between measurements and model output (see Additional Files). Experimental values below the limit of detection (LOD) of the bacterial load per g feces in a given volume of fecal suspension were set to log_10_ CFU/g feces of 2.05 (LOD in the experimental setting C57BL/6J wild type SPF) and run with integrated likelihood. The optimization problem was implemented using the bound-constrained optimization package FMINSEARCHBND in MATLAB 2019 (Mathworks Inc., Massachusetts) and executed on a laptop computer.

## Supporting information

Supplemental File 12 qRTPCR raw data

Supplemental File S13 Data used to calibrate the model

Supplemental File 15 MatLab Script

## Acknowledgements

We thank André Bleich and Marijana Basic from the Institute for Laboratory Animal Science (Hannover Medical School) for providing GF animals. We thank Ulrich Schoppmeier for his great support with statistical analyses. We thank Tanja Späth for technical assistance and all members of the AG Yersinia for their uncomplicated and sustained willingness to support the experiments in a great team effort. Special thanks to Libera Lo Presti for critical reading of the manuscript, deliberate comments and language editing.

## Supplementary Information

**Figure S1.**
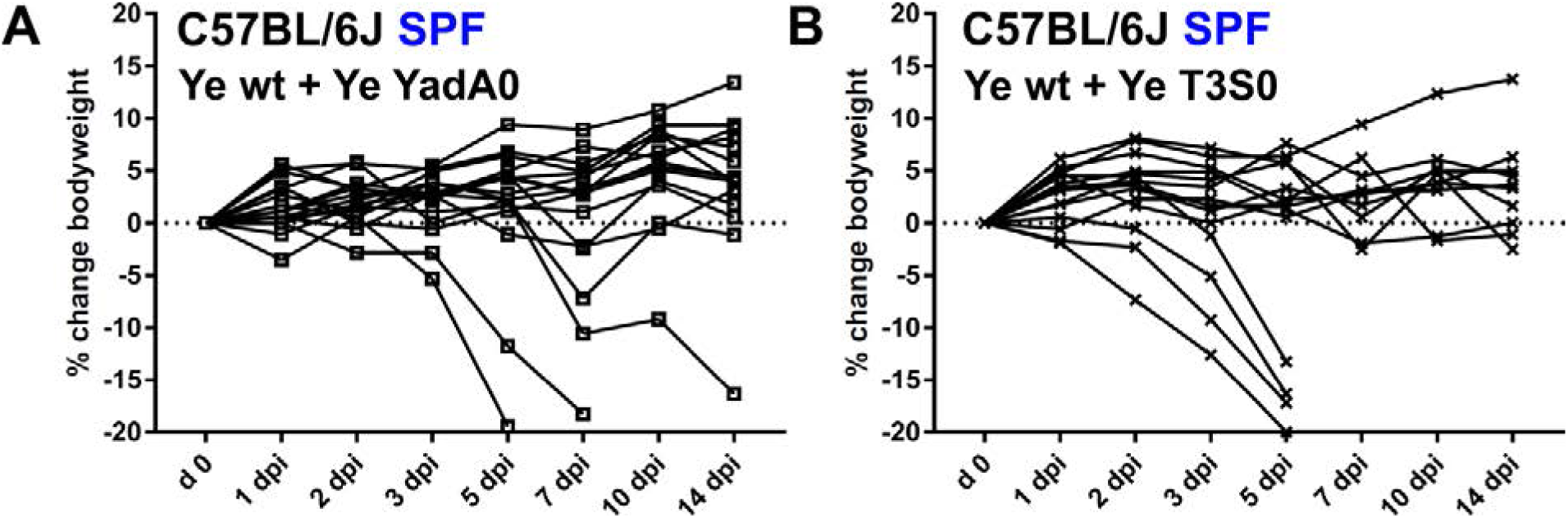
Mouse bodyweight development during Ye coinfection. The bodyweight development of mice during the infection course was monitored as a marker of the severity of the infection. Percent changes compared to initial bodyweight at different time points are illustrated. **(A)** Change of bodyweight of SPF-colonized C57BL/6J mice after 1:1 coinfection with Ye wt and Ye YadA0 and **(B)** with Ye wt and Ye T3S0. As some mice in these coinfections lost significant weight and had to be sacrificed between 5 and 7 dpi, the individual bodyweight developments are shown.

**Figure S2.**
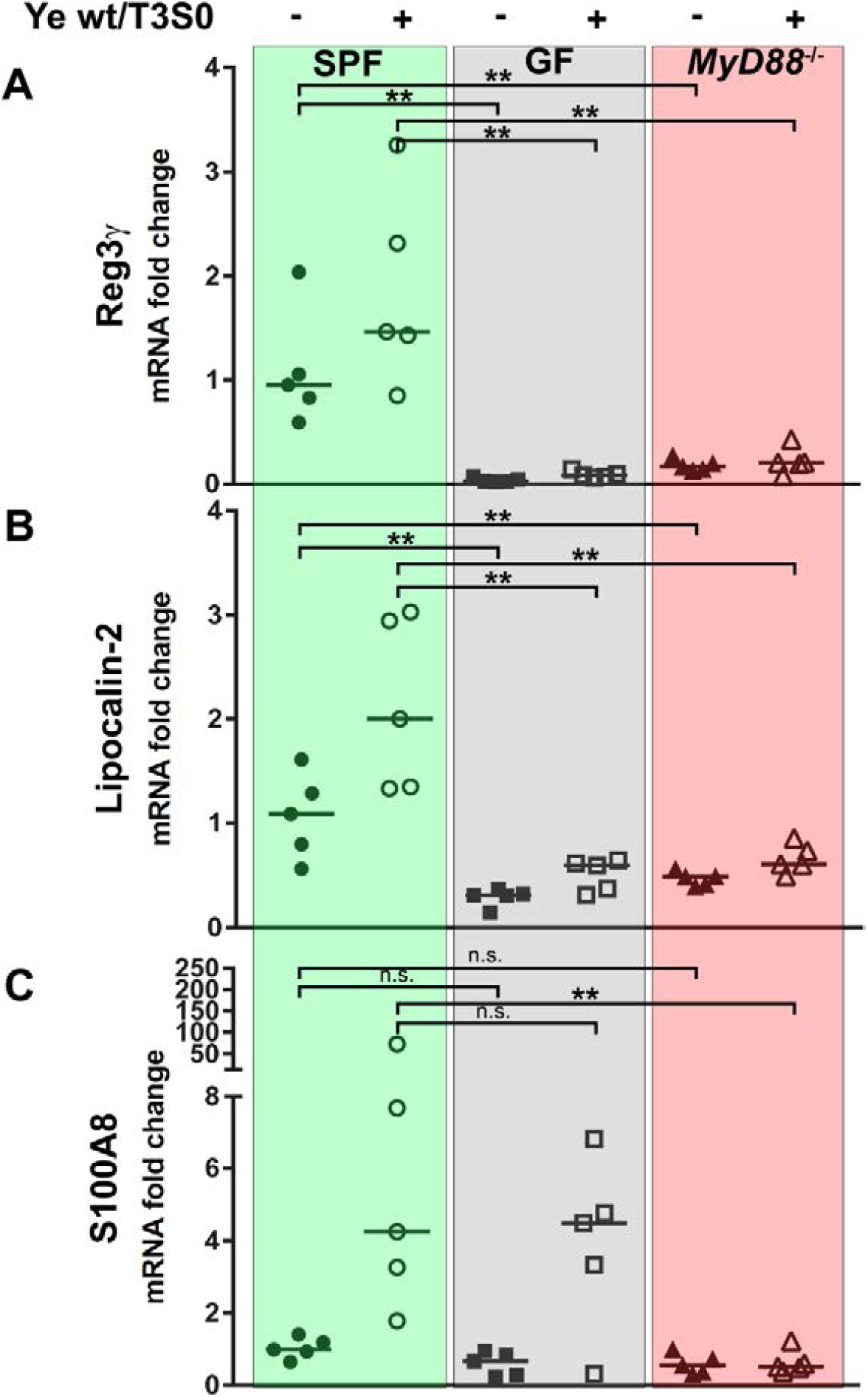
Relative quantification of mRNA levels of Reg3γ, Lipocalin-2 and S100A8 from mucosal scrapings as indicators of intestinal inflammation. Relative expression levels compared to the housekeeping gene beta-glucuronidase were determined by qRT-PCR in mock-infected and mice co-infected for two days with Ye wt/Ye T3S0. **(A)** Basal expression levels and expression levels of Reg3γ following infection of SPF wild type mice, GF animals, and SPF-colonized *MyD88*^-/-^ mice. **(B)** Expression levels of Lipocalin-2 **(C)** Expression levels of S100A8. Statistical significant differences between groups were determined by a nonparametric Mann-Whitney test. ** *P* < 0.01.

**Figure S3.**
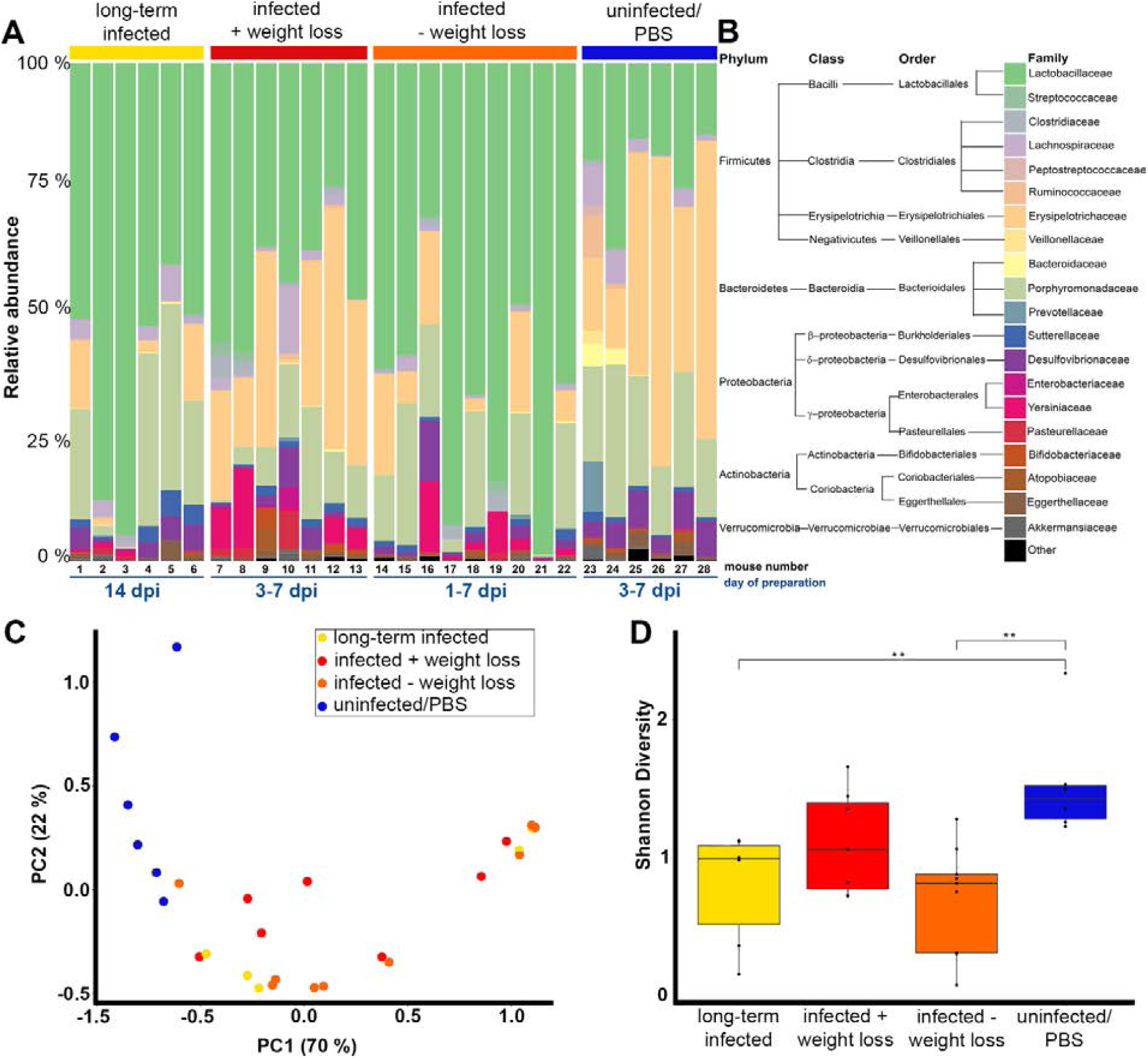
Impact of Ye infection on SI microbiome composition. Microbial composition of SI contents of mice with different infection outcomes, as assessed by 16S rRNA sequencing. (A) Relative abundances of microbiota representatives on the family level. Samples were isolated from the SI after oral Ye wt infection. Relative abundances of families are shown in stacked bar charts for individual animals. Data were grouped according to the observed change of body weight at earlier time points (weight loss after 3-7 days (red), no weight loss between 1 dpi and 7 dpi (orange), or uninfected control group (blue). Long-term infected mice returned to a kind of a steady-state at a late time point of infection and had no signs of sickness anymore (yellow). (B) Taxonomic tree on the phylum-, class-, order- and family-level allowing the assignment of the color code used in (A). (C) Principal component analysis (PC) on Bray-Curtis dissimilarities of the microbial composition of samples. Color code reflects assignments to groups as in (A). (D) Impact of Ye infection on microbial diversity. Shannon diversity of the SI microbiome composition in the different groups of animals. Statistically significant differences were identified using an unpaired Wilcoxon sum rank test. ** P < 0.001.

**Figure S4.**
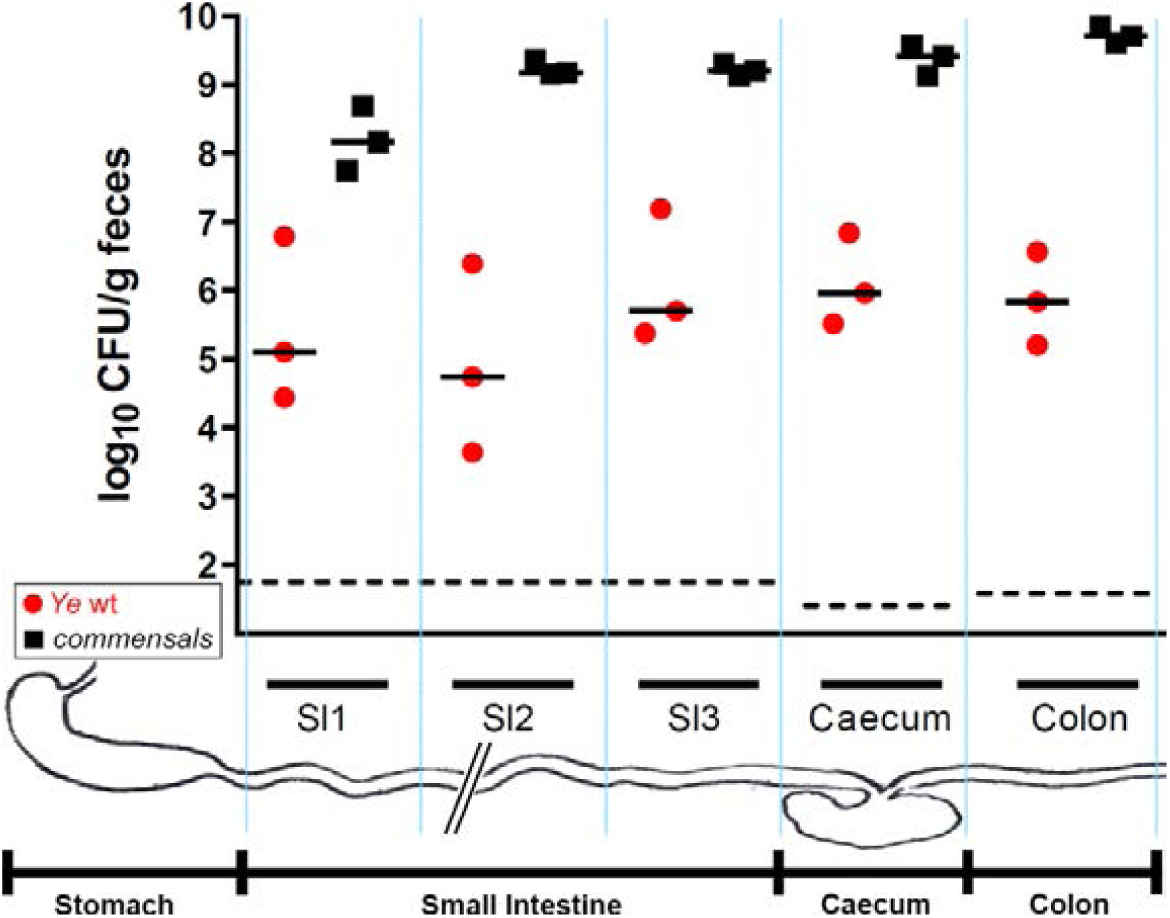
Distribution of Ye and cultivable commensals along the GIT and water content of GI tract sections and fecal pellets. **(A)** At 7 dpi after oral infection of SPF-colonized C57BL/6J mice with the Ye wt strain, the numbers of Ye and cultivable commensals were determined in different compartments of the GIT. The small intestine was dissected and cut into three pieces of equal length (SI1, SI2, SI3). Additionally, the caecum and the colon were dissected. CFUs per gram of fecal content of the three SI pieces and of the caecum and the colon were determined by plating.

**Figure S5.**
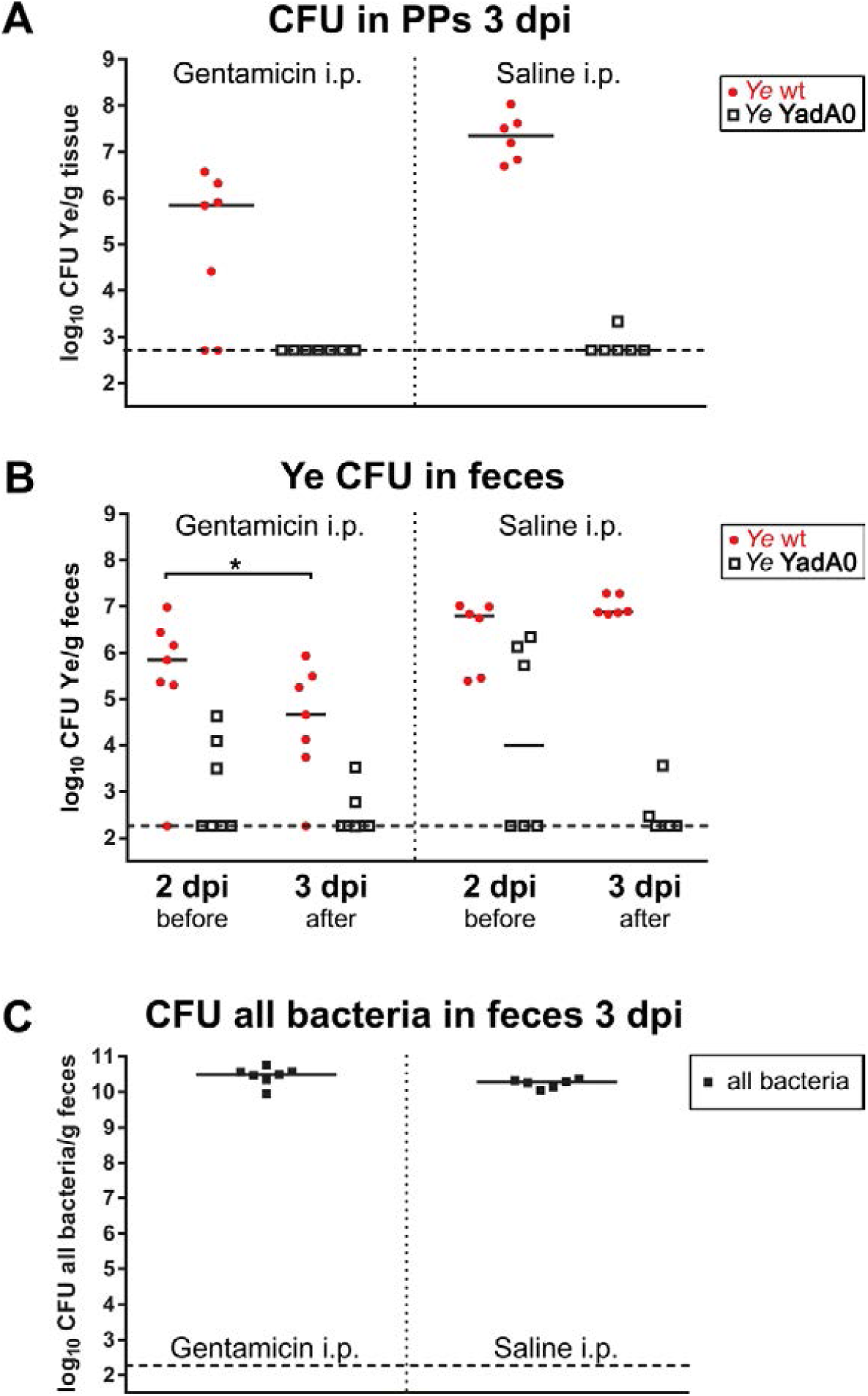
Systemic administration of gentamicin for cleansing of a potential extra-luminal niche colonized by Ye. Mice were orally co-infected with a 1:1 mixture of Ye wt and Ye YadA0 for two days. At this time point, the successful colonization of a potential niche was assumed. One group was then administered intraperitoneally with gentamicin and a control group with saline only. (**A**) Ye wt and Ye YadA0 CFU was determined from Peyer’s patches (PP) on 3 dpi. (**B**) Ye CFU of both strains in feces on days 2 and 3 post-infection. (**C**) The impact of systemic gentamicin treatment on the total CFU at 3 dpi of all cultivable bacteria was addressed by the plating of feces on non-selective agar plates. A paired t-test assessed a statistically significant difference within the Gentamicin treatment group. *P* = 0.494.

**Figure S6.**
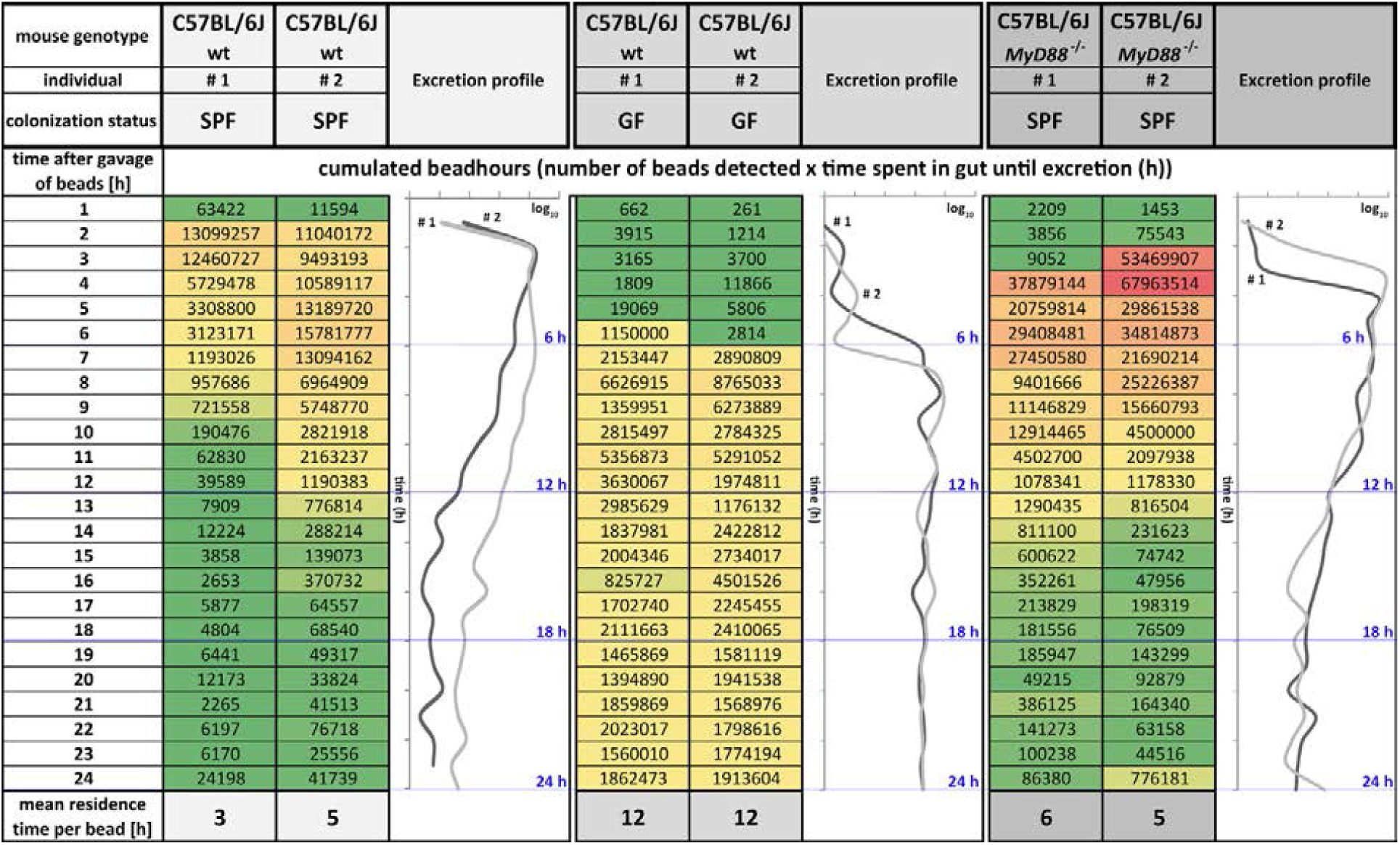
Quantification of gut retention times in SPF-colonized or GF C57BL/6J wild type mice and SPF-colonized *MyD88*^-/-^ animals. Two mice per group were orally challenged with 1·10^9^ fluorescent polystyrene beads plus 5·10^8^ CFU of Ye wt, and feces were collected hourly over 24 h. The number of fluorescent events/g feces at each time point was analyzed by flow cytometry. The cumulated bead-hours were calculated as shown in the heat maps, and the graphs are plotting the log_10_ of cumulated bead-hours for the individual animals. The mean residence time per bead was calculated by dividing the sum of events/g of feces through the total number of bead-hours.

**Figure S7.**
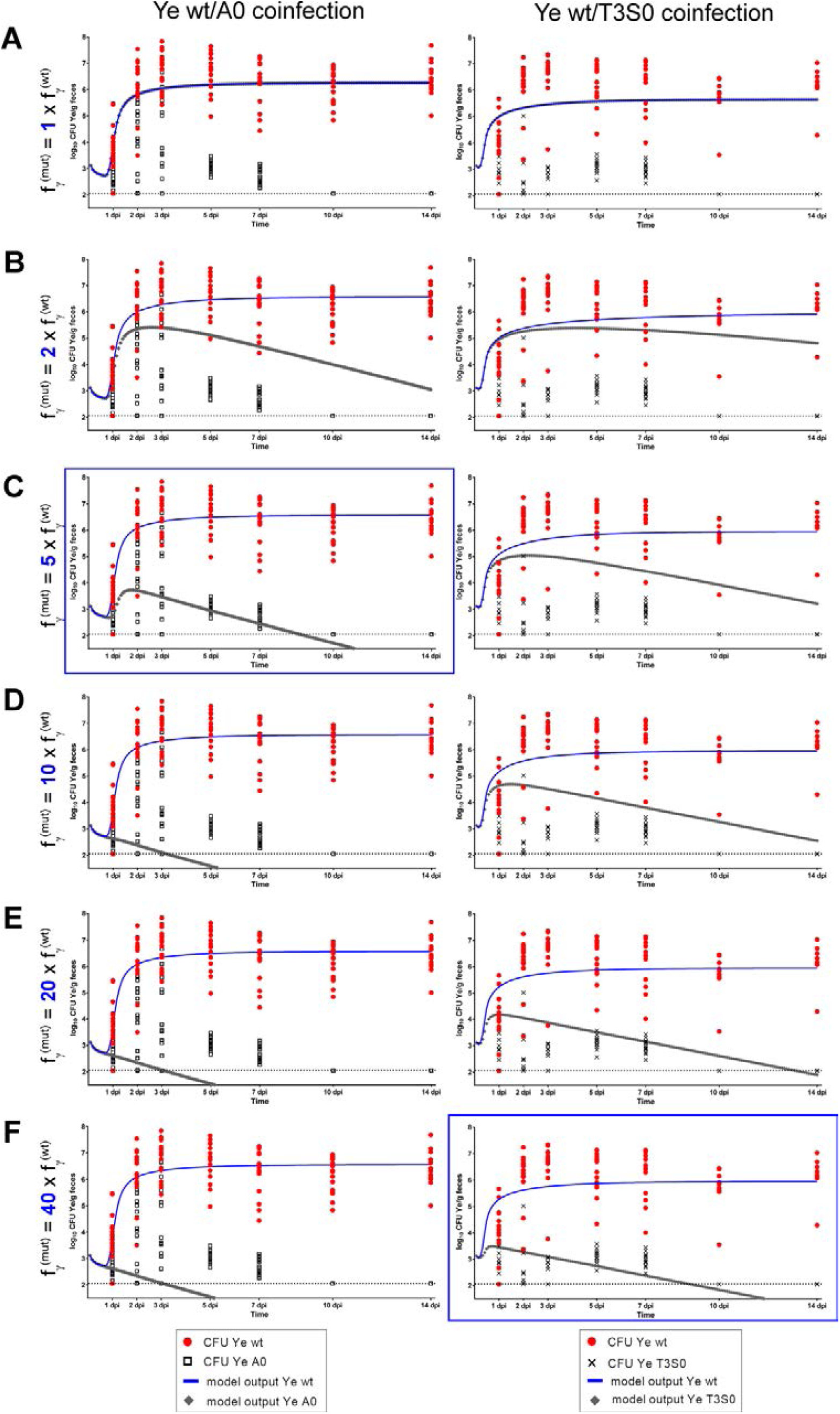
Dynamics of model output when adopting different relations between f_Y_ ^(mut)^ and f_Y_^(wt)^. To visualize the impact of the relative susceptibility to killing by the immune system on population dynamics of the Ye YadA0 (left column) and the Ye T3S0 strain (right column) we plotted curves for f_Y_^(mut)^ adopting values **(A)** equal to that of f_Y_^(wt)^, **(B)** 2x f_Y_ ^(wt)^, **(C)** 5x f_Y_ ^(wt)^, **(D)** 10x f_Y_ ^(wt)^, **(E)** 20x f_Y_ ^(wt)^and **(F)** 40x f_Y_ ^(wt)^ for the respective settings. The relationships that the best matched what our model calculated based on the experimental data are highlighted with a blue frame.

**Supplementary Table S8.** Mean percentage ± SD of water content in sections of the mouse GIT. SI1, SI2, SI3 indicate the respective part of the SI that was analyzed. Please also refer to Figure EV4.

|  | SI1 + ½ SI2 | ½ SI2 + SI3 | Caecum | Colon | Fecal pellet |
| --- | --- | --- | --- | --- | --- |
| **SPF** | 74,64 ± 3,31 | 76,59 ± 3,85 | 75,42 ± 0,35 | 50,69 ± 16,31 | 29,66 ± 2,37 |
| **GF** | 73,11 ± 3,14 | 73,63 ± 0,63 | 78,76 ± 1,79 | 70,07 ± 1,07 | 49,01 ± 3,69 |

**Figure S9.**
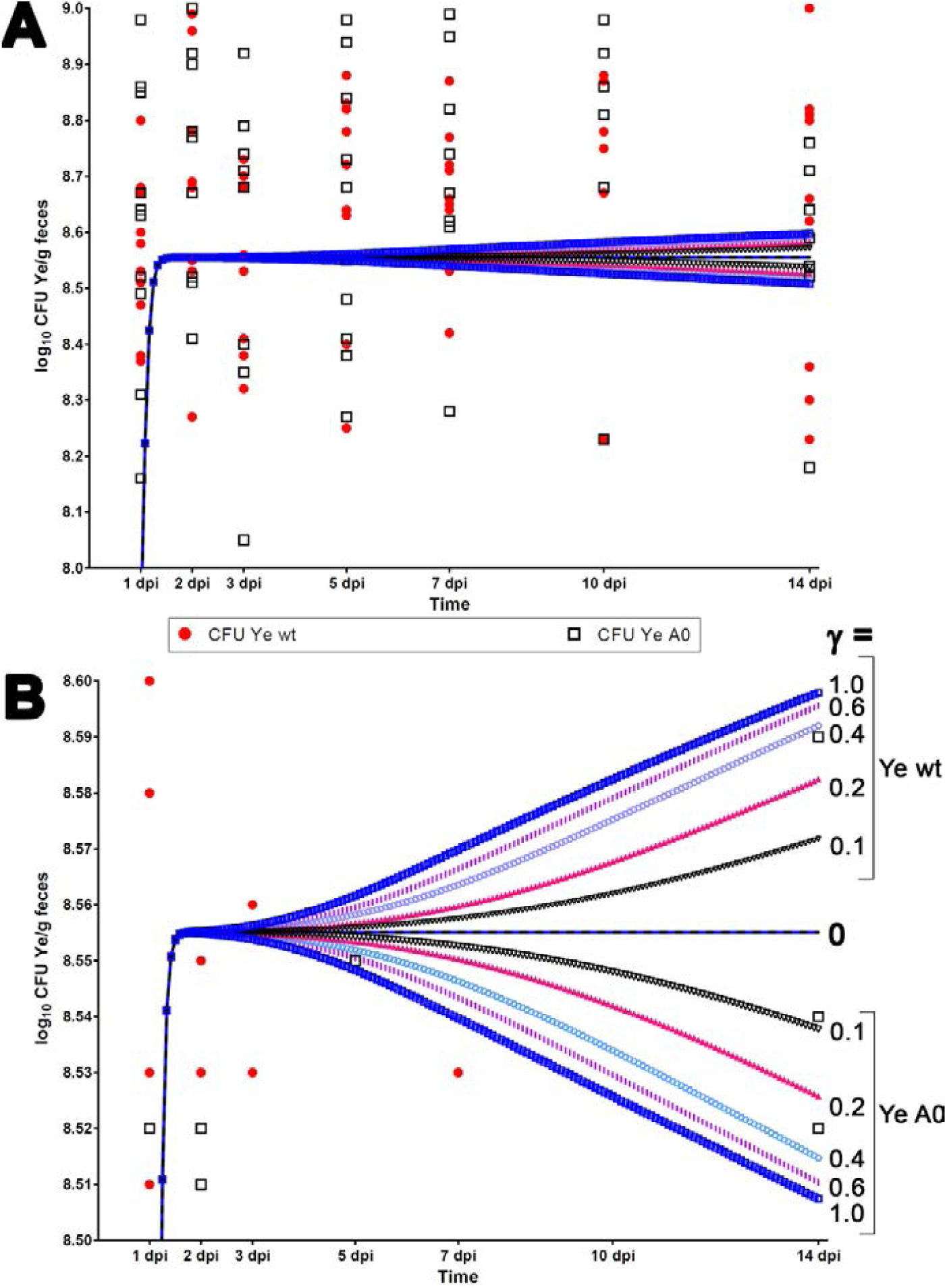
Dynamics of model output in the GF infection setting when adopting different activities of the host immune system. **(A)** Using the same experimental data and parameter set as in Fig. 5A, we calculated the CFU development after the coinfection of GF mice with Ye wt and Ye YadA0 with γ adopting values between 1 (immune system fully active) and 0 (no immune activity). **(B)** For better discrimination of curves, the scale of the y-axis was altered. Curves of the same color and pattern represent one dataset showing the CFU development for Ye wt in the upper half and that of Ye A0 in the lower half of the graph for a value of γ as indicated on the right side. High activity of the immune system correlates with more considerable expansion of the Ye wt strain and decrease of CFU of the Ye A0 strain, but the overall effect of changes in γ is subtle.

**Table S10.**
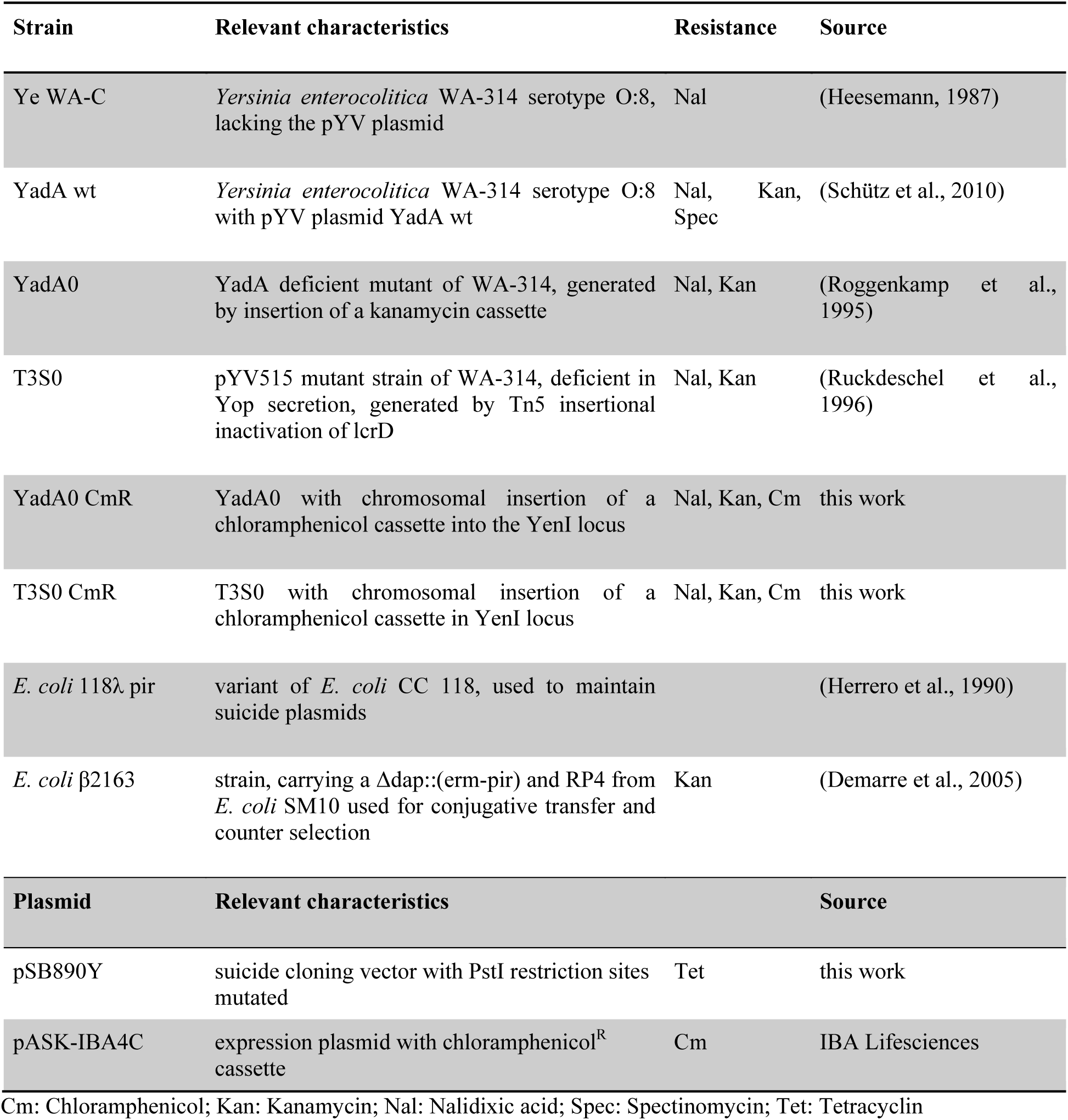
Strains and plasmids used in this study

**Table S11.**
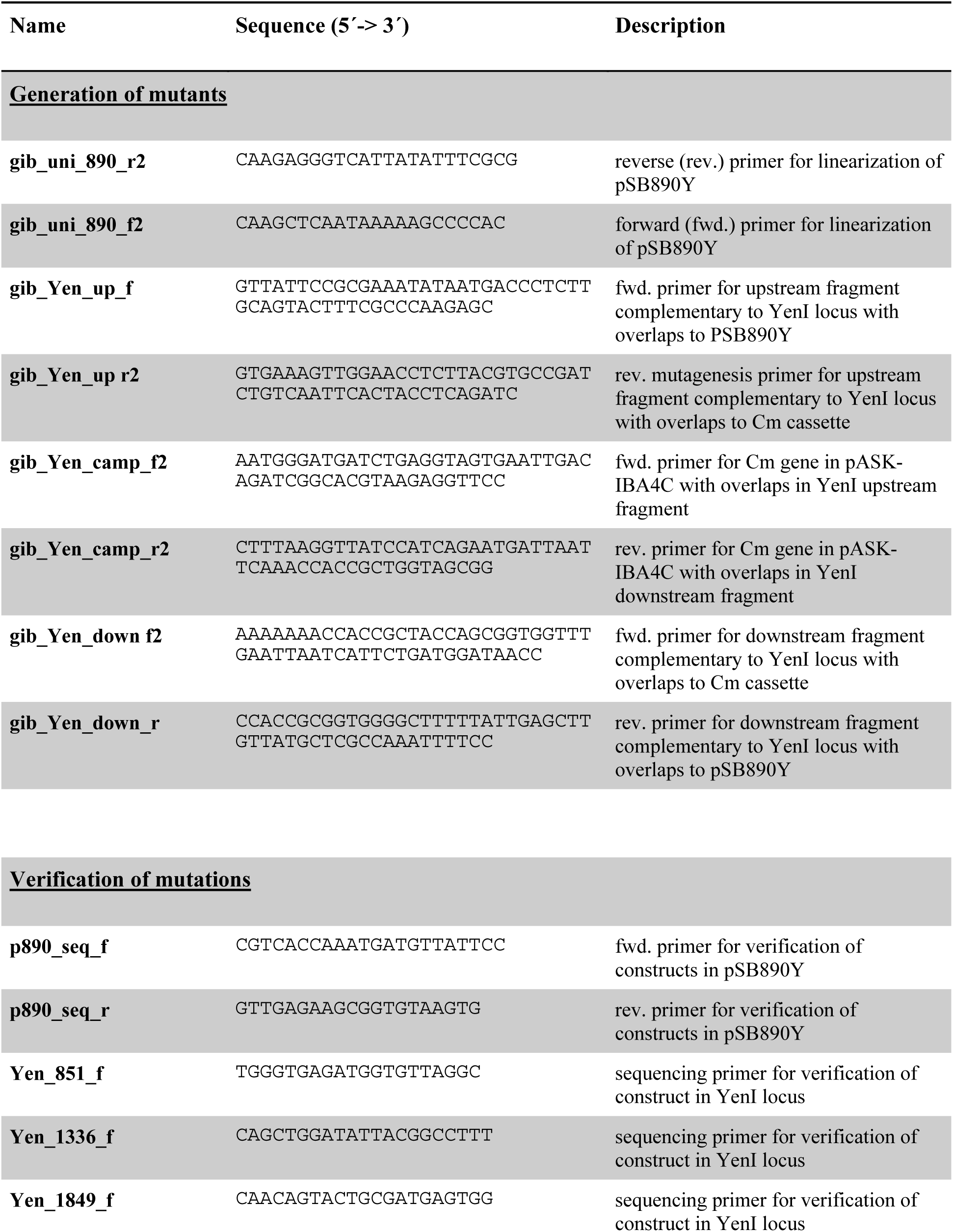
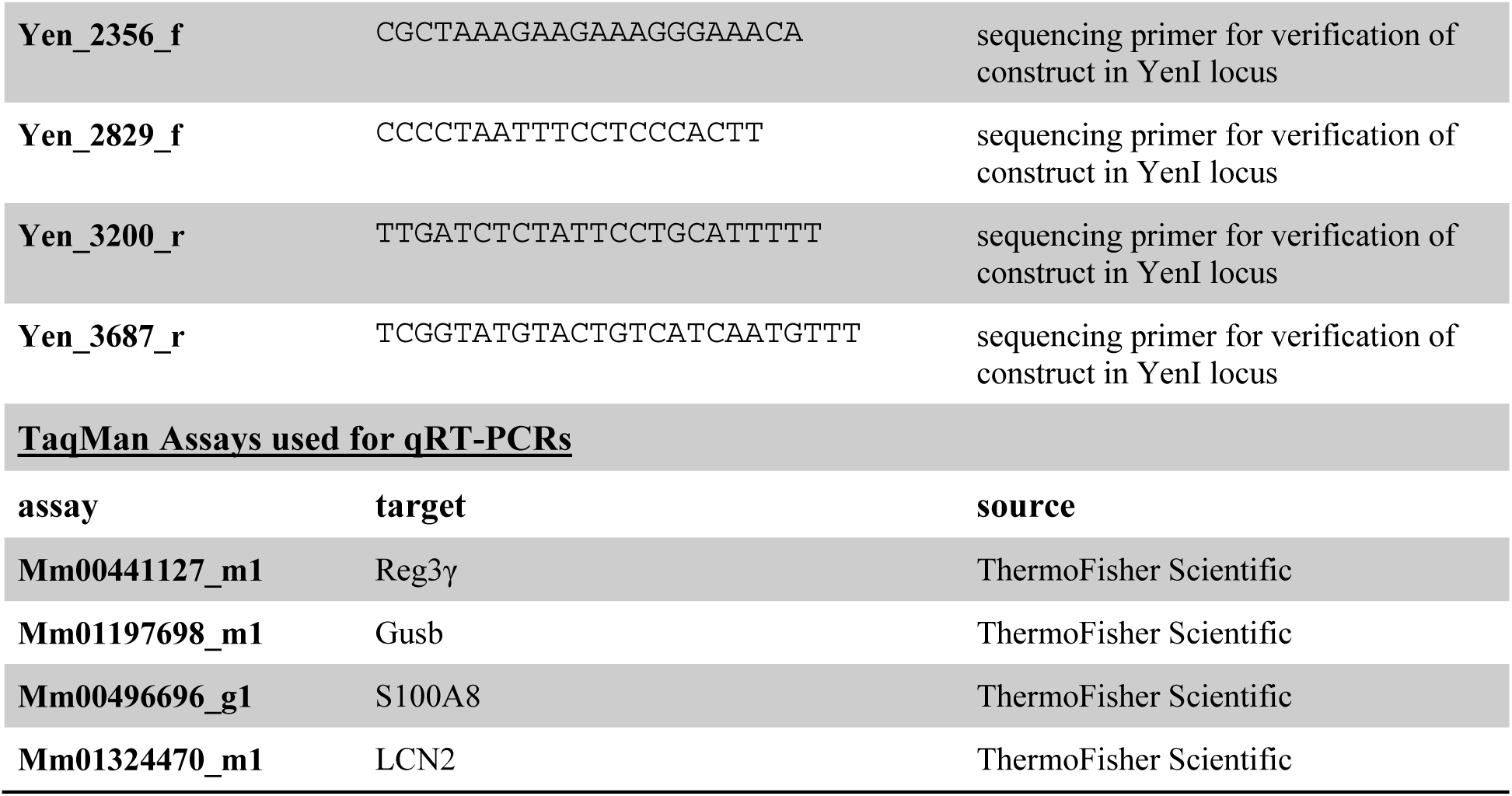
Oligonucleotides used in this study

### Additional files

**S12** qRTPCR raw data.

**S13** The data set used to calibrate the model.

**S14** Computational model in SBML format.

The *Yersinia* colonization model was deposited in BioModels (Chelliah et al., 2015) and assigned the identifier identifiers.org/biomodels.db/MODEL2002070001.

**While under review**, access the models as follows

Please visit https://www.ebi.ac.uk/biomodels/models.

**S15** Matlab script for parameter estimation.

