## Supplemental File 12 qRTPCR raw data for "Model-based prediction of bacterial population dynamics in gastrointestinal infection"

**S12 Table.** qRTPCR raw data.

**Reg3γ**

| **Mouse** | **Treat-ment** | **CP1 Gusb** | **CP2 Gusb** | **Mean CP Gusb** | **Eff. Gusb** | **Eff._Gusb_^Cp(contr.)-Cp(inf.)^** | **CP1 Reg3γ** | **CP2 Reg3γ** | **Mean CP Reg3γ** | **Eff. Reg3γ** | **Eff._Reg3γ_^Cp(contr.)-Cp(inf.)^** | **Fold change** |
| --- | --- | --- | --- | --- | --- | --- | --- | --- | --- | --- | --- | --- |
| **SPF M1** | control | 24,55 | 25,28 | 24,92 | 1,907 | 0,914 | 14,37 | 14,77 | 14,57 | 1,884 | 0,964 | 1,054 |
| **SPF M2** | control | 25,32 | 25,26 | 25,29 | 1,907 | 0,718 | 15,87 | 15,87 | 15,87 | 1,884 | 0,423 | 0,590 |
| **SPF M3** | control | 25,18 | 25,21 | 25,20 | 1,907 | 0,763 | 15,21 | 15,26 | 15,24 | 1,884 | 0,633 | 0,829 |
| **SPF M4** | control | 24,08 | 24,23 | 24,16 | 1,907 | 1,493 | 12,76 | 12,75 | 12,76 | 1,884 | 3,043 | 2,038 |
| **SPF M5** | control | 24,29 | 24,36 | 24,33 | 1,907 | 1,338 | 14,12 | 14,14 | 14,13 | 1,884 | 1,274 | 0,952 |
| **SPF M6** | infected | 23,99 | 23,67 | 23,83 | 1,907 | 1,842 | 13,82 | 13,79 | 13,81 | 1,884 | 1,565 | 0,850 |
| **SPF M7** | infected | 25,20 | 25,34 | 25,27 | 1,907 | 0,727 | 14,40 | 14,51 | 14,46 | 1,884 | 1,037 | 1,426 |
| **SPF M8** | infected | 26,29 | 24,09 | 25,19 | 1,907 | 0,765 | 13,05 | 13,09 | 13,07 | 1,884 | 2,493 | 3,256 |
| **SPF M9** | infected | 24,13 | 24,16 | 24,15 | 1,907 | 1,503 | 13,29 | 13,25 | 13,27 | 1,884 | 2,196 | 1,461 |
| **SPF M10** | infected | 23,54 | 24,20 | 23,87 | 1,907 | 1,795 | 12,25 | 12,28 | 12,27 | 1,884 | 4,151 | 2,313 |
| **GF M1** | control | 27,74 | 26,43 | 27,09 | 1,907 | 0,225 | 21,86 | 21,84 | 21,85 | 1,884 | 0,010 | 0,043 |
| **GF M2** | control | 29,17 | 29,24 | 29,21 | 1,907 | 0,057 | 25,09 | 25,16 | 25,13 | 1,884 | 0,001 | 0,021 |
| **GF M3** | control | 26,99 | 27,10 | 27,05 | 1,907 | 0,231 | 20,99 | 20,98 | 20,99 | 1,884 | 0,017 | 0,072 |
| **GF M4** | control | 25,91 | 26,15 | 26,03 | 1,907 | 0,445 | 21,59 | 21,62 | 21,61 | 1,884 | 0,011 | 0,025 |
| **GF M5** | control | 26,96 | 26,96 | 26,96 | 1,907 | 0,244 | 22,48 | 22,52 | 22,50 | 1,884 | 0,006 | 0,026 |
| **GF M6** | infected | 26,99 | 26,98 | 26,99 | 1,907 | 0,240 | 20,48 | 20,51 | 20,50 | 1,884 | 0,023 | 0,094 |
| **GF M7** | infected | 28,01 | 28,46 | 28,24 | 1,907 | 0,107 | 21,98 | 21,96 | 21,97 | 1,884 | 0,009 | 0,083 |
| **GF M8** | infected | 26,23 | 25,98 | 26,11 | 1,907 | 0,424 | 18,98 | 18,90 | 18,94 | 1,884 | 0,061 | 0,143 |
| **GF M9** | infected | 26,58 | 26,57 | 26,58 | 1,907 | 0,313 | 20,68 | 20,73 | 20,71 | 1,884 | 0,020 | 0,063 |
| **GF M10** | infected | 26,91 | 27,50 | 27,21 | 1,907 | 0,208 | 21,18 | 21,21 | 21,20 | 1,884 | 0,015 | 0,070 |
| **Myd88^-/-^ M1** | control | 25,33 | 25,80 | 25,57 | 1,907 | 0,601 | 17,87 | 17,88 | 17,88 | 1,884 | 0,119 | 0,198 |
| **Myd88^-/-^ M2** | control | 24,92 | 24,91 | 24,92 | 1,907 | 0,914 | 16,73 | 16,70 | 16,72 | 1,884 | 0,248 | 0,271 |
| **Myd88^-/-^ M3** | control | 24,90 | 24,98 | 24,94 | 1,907 | 0,900 | 17,79 | 17,69 | 17,74 | 1,884 | 0,129 | 0,144 |
| **Myd88^-/-^ M4** | control | 25,82 | 25,82 | 25,82 | 1,907 | 0,510 | 18,81 | 18,83 | 18,82 | 1,884 | 0,065 | 0,128 |
| **Myd88^-/-^ M5** | control | 24,68 | 24,73 | 24,71 | 1,907 | 1,047 | 17,13 | 17,39 | 17,26 | 1,884 | 0,175 | 0,168 |
| **Myd88^-/-^ M6** | infected | 27,58 | n.d. | 27,58 | 1,907 | 0,164 | 18,71 | 18,74 | 18,73 | 1,884 | 0,069 | 0,424 |
| **Myd88^-/-^ M7** | infected | 23,96 | 24,01 | 23,99 | 1,907 | 1,666 | 17,69 | 17,72 | 17,71 | 1,884 | 0,132 | 0,079 |
| **Myd88^-/-^ M8** | infected | 24,31 | 24,34 | 24,33 | 1,907 | 1,338 | 16,54 | 16,56 | 16,55 | 1,884 | 0,275 | 0,206 |
| **Myd88^-/-^ M9** | infected | 24,15 | 24,20 | 24,18 | 1,907 | 1,474 | 16,39 | 16,38 | 16,39 | 1,884 | 0,305 | 0,207 |
| **Myd88^-/-^ M10** | infected | 24,10 | 24,05 | 24,08 | 1,907 | 1,572 | 16,32 | 16,46 | 16,39 | 1,884 | 0,304 | 0,194 |

**LCN-2**

| **Mouse** | **Treat-ment** | **CP1 Gusb** | **CP2 Gusb** | **Mean CP Gusb** | **Eff. Gusb** | **Eff._Gusb_^Cp(contr.)-Cp(inf.)^** | **CP1 LCN-2** | **CP2 LCN-2** | **Mean CP LCN-2** | **Eff. LCN-2** | **Eff._LCN-2_^Cp(contr.)-Cp(inf.)^** | **Fold change** |
| --- | --- | --- | --- | --- | --- | --- | --- | --- | --- | --- | --- | --- |
| **SPF M1** | control | 24,37 | 24,44 | 24,41 | 1,820 | 1,095 | 30,58 | 30,67 | 30,63 | 1,591 | 0,870 | 0,795 |
| **SPF M2** | control | 24,86 | 24,84 | 24,85 | 1,820 | 0,839 | 31,90 | 32,00 | 31,95 | 1,591 | 0,470 | 0,561 |
| **SPF M3** | control | 25,21 | 25,16 | 25,19 | 1,820 | 0,686 | 30,60 | 30,60 | 30,60 | 1,591 | 0,880 | 1,283 |
| **SPF M4** | control | 24,07 | 24,12 | 24,10 | 1,820 | 1,318 | 28,70 | 28,71 | 28,71 | 1,591 | 2,122 | 1,610 |
| **SPF M5** | control | 24,25 | 24,24 | 24,25 | 1,820 | 1,205 | 29,68 | 29,81 | 29,75 | 1,591 | 1,309 | 1,087 |
| **SPF M6** | infected | 23,82 | 23,85 | 23,84 | 1,820 | 1,540 | 28,82 | 28,75 | 28,79 | 1,591 | 2,044 | 1,328 |
| **SPF M7** | infected | 25,08 | 25,14 | 25,11 | 1,820 | 0,718 | 30,32 | 30,48 | 30,40 | 1,591 | 0,966 | 1,346 |
| **SPF M8** | infected | 24,08 | 24,07 | 24,08 | 1,820 | 1,334 | 27,37 | 27,39 | 27,38 | 1,591 | 3,926 | 2,943 |
| **SPF M9** | infected | 24,06 | 24,12 | 24,09 | 1,820 | 1,322 | 28,22 | 28,24 | 28,23 | 1,591 | 2,645 | 2,001 |
| **SPF M10** | infected | 23,52 | 23,53 | 23,53 | 1,820 | 1,854 | 26,59 | 26,63 | 26,61 | 1,591 | 5,613 | 3,027 |
| **GF M1** | control | 26,3 | 26,46 | 26,38 | 1,820 | 0,335 | 35,32 | 35,08 | 35,20 | 1,591 | 0,104 | 0,310 |
| **GF M2** | control | 28,97 | 28,91 | 28,94 | 1,820 | 0,072 | 38,13 | 38,97 | 38,55 | 1,591 | 0,022 | 0,303 |
| **GF M3** | control | 26,90 | 27,01 | 26,96 | 1,820 | 0,238 | 37,59 | 37,61 | 37,60 | 1,591 | 0,034 | 0,143 |
| **GF M4** | control | 26,10 | 26,05 | 26,08 | 1,820 | 0,403 | 34,55 | 34,35 | 34,45 | 1,591 | 0,147 | 0,366 |
| **GF M5** | control | 26,89 | 26,84 | 26,87 | 1,820 | 0,251 | 35,63 | 35,88 | 35,76 | 1,591 | 0,080 | 0,320 |
| **GF M6** | infected | 26,81 | 26,84 | 26,83 | 1,820 | 0,257 | 35,73 | 35,83 | 35,78 | 1,591 | 0,079 | 0,309 |
| **GF M7** | infected | 27,78 | 27,79 | 27,79 | 1,820 | 0,145 | 35,75 | 35,48 | 35,62 | 1,591 | 0,086 | 0,593 |
| **GF M8** | infected | 25,99 | 25,95 | 25,97 | 1,820 | 0,429 | 33,13 | 33,06 | 33,10 | 1,591 | 0,276 | 0,644 |
| **GF M9** | infected | 26,46 | 26,35 | 26,41 | 1,820 | 0,330 | 33,62 | 33,89 | 33,76 | 1,591 | 0,203 | 0,615 |
| **GF M10** | infected | 26,90 | 26,97 | 26,94 | 1,820 | 0,241 | 35,49 | 35,58 | 35,54 | 1,591 | 0,089 | 0,370 |
| **Myd88^-/-^ M1** | control | 24,64 | 24,63 | 24,64 | 1,820 | 0,954 | 32,01 | 32,85 | 32,43 | 1,591 | 0,376 | 0,394 |
| **Myd88^-/-^ M2** | control | 24,36 | 24,35 | 24,36 | 1,820 | 1,128 | 31,63 | 31,60 | 31,62 | 1,591 | 0,549 | 0,487 |
| **Myd88^-/-^ M3** | control | 24,79 | 24,77 | 24,78 | 1,820 | 0,874 | 32,49 | 32,52 | 32,51 | 1,591 | 0,363 | 0,416 |
| **Myd88^-/-^ M4** | control | 25,92 | 25,95 | 25,94 | 1,820 | 0,438 | 33,71 | 33,57 | 33,64 | 1,591 | 0,215 | 0,490 |
| **Myd88^-/-^ M5** | control | 23,91 | 23,89 | 23,90 | 1,820 | 1,481 | 30,67 | 30,79 | 30,73 | 1,591 | 0,829 | 0,559 |
| **Myd88^-/-^ M6** | infected | 25,17 | 25,19 | 25,18 | 1,820 | 0,688 | 32,59 | 32,72 | 32,66 | 1,591 | 0,339 | 0,492 |
| **Myd88^-/-^ M7** | infected | 23,87 | 23,97 | 23,92 | 1,820 | 1,464 | 30,58 | 30,63 | 30,61 | 1,591 | 0,878 | 0,600 |
| **Myd88^-/-^ M8** | infected | 24,08 | 24,15 | 24,12 | 1,820 | 1,302 | 30,48 | 30,38 | 30,43 | 1,591 | 0,952 | 0,731 |
| **Myd88^-/-^ M9** | infected | 24,30 | 24,48 | 24,39 | 1,820 | 1,105 | 30,33 | 30,58 | 30,46 | 1,591 | 0,941 | 0,852 |
| **Myd88^-/-^ M10** | infected | 24,03 | 24,04 | 24,04 | 1,820 | 1,366 | 30,73 | 30,73 | 30,73 | 1,591 | 0,829 | 0,606 |

**S100A8**

| **Mouse** | **Treat-ment** | **CP1 Gusb** | **CP2 Gusb** | **Mean CP Gusb** | **Eff. Gusb** | **Eff._Gusb_^Cp(contr.)-Cp(inf.)^** | **CP1 S100A8** | **CP2 S100A8** | **Mean CP S100A8** | **Eff. S100A8** | **Eff._S100A8_^Cp(contr.)-Cp(inf.)^** | **Fold change** |
| --- | --- | --- | --- | --- | --- | --- | --- | --- | --- | --- | --- | --- |
| **SPF M1** | control | 24,44 | 24,46 | 24,45 | 1,886 | 1,059 | 33,80 | 33,53 | 33,67 | 1,678 | 1,050 | 0,991 |
| **SPF M2** | control | 24,83 | 24,9 | 24,87 | 1,886 | 0,814 | 33,63 | 33,37 | 33,50 | 1,678 | 1,143 | 1,404 |
| **SPF M3** | control | 25,15 | 25,11 | 25,13 | 1,886 | 0,688 | 35,10 | 35,53 | 35,32 | 1,678 | 0,447 | 0,649 |
| **SPF M4** | control | 24,03 | 24,1 | 24,07 | 1,886 | 1,353 | 33,28 | 33,33 | 33,31 | 1,678 | 1,265 | 0,935 |
| **SPF M5** | control | 24,23 | 24,16 | 24,20 | 1,886 | 1,245 | 32,73 | 33,29 | 33,01 | 1,678 | 1,474 | 1,183 |
| **SPF M6** | infected | 23,80 | 23,77 | 23,79 | 1,886 | 1,616 | 30,62 | 30,47 | 30,55 | 1,678 | 5,278 | 3,267 |
| **SPF M7** | infected | 25,29 | 25,15 | 25,22 | 1,886 | 0,650 | 33,48 | 33,47 | 33,48 | 1,678 | 1,158 | 1,782 |
| **SPF M8** | infected | 24,24 | 24,24 | 24,24 | 1,886 | 1,210 | 25,07 | 25,16 | 25,12 | 1,678 | 87,721 | 72,471 |
| **SPF M9** | infected | 24,15 | 24,15 | 24,15 | 1,886 | 1,282 | 30,49 | 30,48 | 30,49 | 1,678 | 5,445 | 4,248 |
| **SPF M10** | infected | 23,72 | 23,5 | 23,61 | 1,886 | 1,805 | 28,66 | 28,70 | 28,68 | 1,678 | 13,859 | 7,677 |
| **GF M1** | control | 26,46 | 26,34 | 26,40 | 1,886 | 0,307 | 36,74 | 36,04 | 36,39 | 1,678 | 0,256 | 0,833 |
| **GF M2** | control | 29,29 | 28,98 | 29,14 | 1,886 | 0,054 | 39,85 | 39,09 | 39,47 | 1,678 | 0,052 | 0,960 |
| **GF M3** | control | 26,85 | 26,85 | 26,85 | 1,886 | 0,231 | 39,10 | 38,97 | 39,04 | 1,678 | 0,065 | 0,282 |
| **GF M4** | control | 26,14 | 26,08 | 26,11 | 1,886 | 0,370 | 38,36 | 38,52 | 38,44 | 1,678 | 0,089 | 0,240 |
| **GF M5** | control | 26,90 | 26,92 | 26,91 | 1,886 | 0,222 | 37,14 | 37,75 | 37,45 | 1,678 | 0,148 | 0,667 |
| **GF M6** | infected | 27,81 | 28,07 | 27,94 | 1,886 | 0,116 | 34,30 | 34,14 | 34,22 | 1,678 | 0,788 | 6,807 |
| **GF M7** | infected | 27,46 | 26,97 | 27,22 | 1,886 | 0,183 | 33,89 | 34,15 | 34,02 | 1,678 | 0,874 | 4,766 |
| **GF M8** | infected | 25,97 | 26,03 | 26,00 | 1,886 | 0,396 | 32,65 | 32,64 | 32,65 | 1,678 | 1,780 | 4,492 |
| **GF M9** | infected | 26,44 | 26,64 | 26,54 | 1,886 | 0,281 | 33,90 | 33,86 | 33,88 | 1,678 | 0,939 | 3,339 |
| **GF M10** | infected | 27,10 | 27,17 | 27,14 | 1,886 | 0,193 | 38,92 | 39,44 | 39,18 | 1,678 | 0,060 | 0,313 |
| **Myd88^-/-^ M1** | control | 24,51 | 24,56 | 24,54 | 1,886 | 1,004 | 35,84 | 35,38 | 35,61 | 1,678 | 0,384 | 0,382 |
| **Myd88^-/-^ M2** | control | 24,53 | 24,44 | 24,49 | 1,886 | 1,036 | 33,54 | 33,86 | 33,70 | 1,678 | 1,031 | 0,995 |
| **Myd88^-/-^ M3** | control | 24,7 | 24,82 | 24,76 | 1,886 | 0,870 | 35,16 | 35,14 | 35,15 | 1,678 | 0,487 | 0,559 |
| **Myd88^-/-^ M4** | control | 26,00 | 26,18 | 26,09 | 1,886 | 0,374 | 36,62 | 35,91 | 36,27 | 1,678 | 0,273 | 0,730 |
| **Myd88^-/-^ M5** | control | 23,84 | 23,90 | 23,87 | 1,886 | 1,531 | 34,77 | 35,55 | 35,16 | 1,678 | 0,484 | 0,316 |
| **Myd88^-/-^ M6** | infected | 25,15 | 25,17 | 25,16 | 1,886 | 0,675 | 35,86 | 35,14 | 35,50 | 1,678 | 0,406 | 0,601 |
| **Myd88^-/-^ M7** | infected | 24,06 | 24,03 | 24,05 | 1,886 | 1,370 | 34,73 | 35,32 | 35,03 | 1,678 | 0,519 | 0,379 |
| **Myd88^-/-^ M8** | infected | 24,01 | 24,03 | 24,02 | 1,886 | 1,392 | 32,78 | 32,68 | 32,73 | 1,678 | 1,703 | 1,224 |
| **Myd88^-/-^ M9** | infected | 24,55 | 24,17 | 24,36 | 1,886 | 1,122 | 34,81 | 34,99 | 34,90 | 1,678 | 0,554 | 0,494 |
| **Myd88^-/-^ M10** | infected | 24,44 | 24,34 | 24,39 | 1,886 | 1,101 | 34,77 | 34,96 | 34,87 | 1,678 | 0,564 | 0,513 |
