## Supplemental File S13 Data used to calibrate the model for "Model-based prediction of bacterial population dynamics in gastrointestinal infection"

**S13 Table.** Data set used to calibrate the model.

SPF wildtype/A0 coinfection

| **Time (hours post infection)** | **log10 CFU/g feces of Ye wildtype in lumen** | **log10 CFU/g feces of Ye YadA0 in lumen** |
| --- | --- | --- |
| 24 | 3,08 | 3,22 |
| 24 | 3,39 | 2,41 |
| 24 | 2,05 | 2,16 |
| 24 | 2,05 | 2,11 |
| 24 | 4,18 | 2,37 |
| 24 | 3,56 | 2,44 |
| 24 | 4,21 | 2,49 |
| 24 | 3,59 | 2,73 |
| 24 | 3,72 | 3,22 |
| 24 | 2,05 | 2,60 |
| 24 | 3,28 | 2,86 |
| 24 | 4,16 | 3,44 |
| 24 | 3,89 | 2,97 |
| 24 | 5,47 | 3,85 |
| 24 | 4,02 | 3,35 |
| 24 | 4,65 | 3,88 |
| 24 | 5,44 | 3,16 |
| 48 | 6,21 | 5,17 |
| 48 | 6,76 | 5,79 |
| 48 | 5,73 | 2,05 |
| 48 | 5,84 | 2,05 |
| 48 | 7,04 | 4,78 |
| 48 | 6,61 | 4,58 |
| 48 | 5,85 | 5,48 |
| 48 | 6,64 | 3,86 |
| 48 | 4,53 | 2,05 |
| 48 | 5,90 | 4,15 |
| 48 | 3,50 | 2,52 |
| 48 | 6,72 | 5,64 |
| 48 | 6,06 | 2,07 |
| 48 | 7,10 | 5,02 |
| 48 | 6,85 | 4,05 |
| 48 | 7,54 | 2,79 |
| 48 | 6,19 | 2,05 |
| 72 | 6,44 | 5,13 |
| 72 | 7,16 | 6,10 |
| 72 | 6,37 | 3,53 |
| 72 | 6,77 | 3,19 |
| 72 | 7,50 | 3,55 |
| 72 | 6,93 | 2,61 |
| 72 | 7,23 | 5,58 |
| 72 | 7,12 | 3,09 |
| 72 | 5,80 | 3,79 |
| 72 | 7,01 | 3,59 |
| 72 | 5,43 | 2,05 |
| 72 | 7,13 | 2,05 |
| 72 | 5,72 | 2,05 |
| 72 | 7,84 | 5,02 |
| 72 | 7,60 | 6,66 |
| 72 | 6,35 | 2,05 |
| 72 | 5,91 | 2,05 |
| 120 | 7,18 | 3,07 |
| 120 | 6,99 | 3,00 |
| 120 | 6,76 | 3,03 |
| 120 | 6,74 | 3,29 |
| 120 | 6,11 | 2,66 |
| 120 | 6,34 | 2,87 |
| 120 | 7,65 | 2,85 |
| 120 | 6,56 | 3,34 |
| 120 | 5,81 | 3,02 |
| 120 | 7,35 | 3,29 |
| 120 | 4,98 | 3,28 |
| 120 | 6,77 | 3,48 |
| 120 | 6,85 | 3,13 |
| 120 | 7,51 | 3,01 |
| 120 | 7,38 | 2,79 |
| 120 | 6,12 | 3,19 |
| 120 | 5,62 | 2,96 |
| 168 | 5,56 | 2,27 |
| 168 | 6,22 | 2,98 |
| 168 | 6,46 | 2,36 |
| 168 | 6,32 | 2,96 |
| 168 | 7,24 | 2,45 |
| 168 | 6,96 | 2,87 |
| 168 | 6,32 | 2,62 |
| 168 | 7,00 | 2,39 |
| 168 | 5,07 | 3,09 |
| 168 | 7,17 | 3,05 |
| 168 | 4,44 | 2,86 |
| 168 | 6,29 | 2,68 |
| 168 | 6,60 | 3,17 |
| 168 | 7,26 | 2,93 |
| 168 | 7,24 | 2,54 |
| 168 | 4,84 | 2,98 |
| 240 | 5,60 | 2,05 |
| 240 | 6,33 | 2,05 |
| 240 | 6,62 | 2,05 |
| 240 | 5,47 | 2,05 |
| 240 | 6,55 | 2,05 |
| 240 | 5,91 | 2,05 |
| 240 | 6,51 | 2,05 |
| 240 | 6,30 | 2,05 |
| 240 | 6,94 | 2,05 |
| 240 | 5,12 | 2,05 |
| 240 | 5,53 | 2,05 |
| 240 | 6,77 | 2,05 |
| 240 | 6,11 | 2,05 |
| 240 | 6,85 | 2,05 |
| 240 | 4,84 | 2,05 |
| 336 | 6,42 | 2,05 |
| 336 | 6,65 | 2,05 |
| 336 | 7,16 | 2,05 |
| 336 | 5,88 | 2,05 |
| 336 | 6,01 | 2,05 |
| 336 | 6,31 | 2,05 |
| 336 | 7,04 | 2,05 |
| 336 | 6,64 | 2,05 |
| 336 | 7,68 | 2,05 |
| 336 | 5,01 | 2,05 |
| 336 | 6,43 | 2,05 |
| 336 | 6,07 | 2,05 |
| 336 | 6,39 | 2,05 |
| 336 | 6,16 | 2,05 |
| 336 | 6,75 | 2,05 |

SPF wildtype/A0 coinfection median

| **Time (hours post infection)** | **median of log10 CFU/g feces of Ye wildtype in lumen** | **median of log10 CFU/g feces of Ye YadA0 in lumen** |
| --- | --- | --- |
| 24 | 3,72 | 2,86 |
| 48 | 6,21 | 4,05 |
| 72 | 6,93 | 3,53 |
| 120 | 6,76 | 3,03 |
| 168 | 6,39 | 2,87 |
| 240 | 6,30 | 2,05 |
| 336 | 6,42 | 2,05 |

SPF wildtype/T3S0 coinfection

| **Time (hours post infection)** | **log10 CFU/g feces of Ye wildtype in lumen** | **log10 CFU/g feces of Ye T3S0 in lumen** |
| --- | --- | --- |
| 24 | 2,66 | 2,66 |
| 24 | 2,05 | 2,05 |
| 24 | 5,67 | 4,01 |
| 24 | 4,14 | 2,84 |
| 24 | 4,45 | 2,87 |
| 24 | 3,61 | 2,46 |
| 24 | 4,26 | 2,05 |
| 24 | 3,91 | 2,05 |
| 24 | 5,35 | 3,47 |
| 24 | 4,00 | 2,05 |
| 24 | 4,78 | 3,00 |
| 24 | 3,71 | 2,05 |
| 24 | 3,70 | 3,22 |
| 24 | 4,75 | 2,05 |
| 48 | 4,56 | 2,05 |
| 48 | 3,37 | 2,05 |
| 48 | 6,15 | 2,05 |
| 48 | 6,84 | 2,05 |
| 48 | 6,58 | 2,11 |
| 48 | 6,12 | 2,05 |
| 48 | 5,94 | 2,05 |
| 48 | 6,17 | 3,23 |
| 48 | 7,24 | 5,02 |
| 48 | 6,53 | 2,50 |
| 48 | 6,26 | 2,45 |
| 48 | 6,75 | 2,21 |
| 48 | 6,37 | 2,05 |
| 48 | 6,65 | 2,05 |
| 72 | 6,91 | 2,05 |
| 72 | 3,76 | 2,05 |
| 72 | 7,06 | 2,05 |
| 72 | 6,63 | 2,73 |
| 72 | 6,67 | 3,08 |
| 72 | 6,35 | 2,05 |
| 72 | 6,70 | 2,05 |
| 72 | 6,08 | 2,88 |
| 72 | 7,35 | 2,86 |
| 72 | 6,83 | 2,63 |
| 72 | 6,73 | 2,05 |
| 72 | 6,78 | 2,05 |
| 72 | 7,04 | 3,04 |
| 72 | 7,31 | 2,05 |
| 120 | 6,30 | 2,85 |
| 120 | 4,34 | 2,57 |
| 120 | 7,13 | 3,30 |
| 120 | 5,31 | 3,23 |
| 120 | 5,78 | 3,14 |
| 120 | 5,90 | 3,05 |
| 120 | 6,99 | 3,03 |
| 120 | 6,77 | 3,57 |
| 120 | 6,70 | 3,21 |
| 120 | 5,73 | 3,21 |
| 120 | 7,14 | 3,41 |
| 120 | 6,90 | 3,03 |
| 120 | 6,10 | 2,94 |
| 120 | 6,80 | 3,40 |
| 168 | 5,24 | 3,31 |
| 168 | 4,01 | 2,46 |
| 168 | 6,56 | 2,89 |
| 168 | 6,78 | 2,98 |
| 168 | 7,07 | 3,13 |
| 168 | 6,92 | 2,69 |
| 168 | 5,51 | 2,91 |
| 168 | 6,44 | 3,43 |
| 168 | 6,29 | 3,08 |
| 168 | 7,13 | 2,86 |
| 168 | 6,41 | 2,74 |
| 168 | 4,95 | 3,08 |
| 168 | 7,13 | 3,17 |
| 240 | 5,85 | 2,05 |
| 240 | 3,54 | 2,05 |
| 240 | 5,71 | 2,05 |
| 240 | 5,72 | 2,05 |
| 240 | 5,56 | 2,05 |
| 240 | 5,62 | 2,05 |
| 240 | 6,42 | 2,05 |
| 240 | 5,91 | 2,05 |
| 240 | 6,45 | 2,05 |
| 240 | 6,15 | 2,05 |
| 336 | 6,08 | 2,05 |
| 336 | 4,29 | 2,05 |
| 336 | 7,03 | 2,05 |
| 336 | 6,08 | 2,05 |
| 336 | 6,32 | 2,05 |
| 336 | 6,69 | 2,05 |
| 336 | 6,50 | 2,05 |
| 336 | 6,18 | 2,05 |
| 336 | 6,14 | 2,05 |
| 336 | 6,32 | 2,05 |

SPF wildtype/T3S0 coinfection median

| **Time (hours post infection)** | **median of log10 CFU/g feces of Ye wildtype in lumen** | **median of log10 CFU/g feces of Ye T3S0 in lumen** |
| --- | --- | --- |
| 24 | 4,07 | 2,56 |
| 48 | 6,32 | 2,05 |
| 72 | 6,76 | 2,05 |
| 120 | 6,50 | 3,18 |
| 168 | 6,44 | 2,98 |
| 240 | 5,79 | 2,05 |
| 336 | 6,25 | 2,05 |

GF wildtype/A0 coinfection

| **Time (hours post infection)** | **log10 CFU/g feces of Ye wildtype in lumen** | **log10 CFU/g feces of Ye YadA0 in lumen** |
| --- | --- | --- |
| 24 | 8,47 | 8,52 |
| 24 | 8,58 | 8,67 |
| 24 | 8,51 | 8,86 |
| 24 | 8,37 | 8,49 |
| 24 | 8,60 | 8,64 |
| 24 | 8,80 | 8,63 |
| 24 | 8,68 | 8,16 |
| 24 | 8,67 | 8,85 |
| 24 | 8,38 | 8,31 |
| 24 | 8,53 | 8,98 |
| 48 | 8,96 | 8,77 |
| 48 | 8,78 | 8,78 |
| 48 | 8,69 | 8,90 |
| 48 | 8,96 | 9,18 |
| 48 | 8,53 | 8,52 |
| 48 | 8,99 | 9,00 |
| 48 | 8,68 | 8,92 |
| 48 | 9,07 | 8,41 |
| 48 | 8,27 | 8,67 |
| 48 | 8,55 | 8,51 |
| 72 | 8,38 | 8,79 |
| 72 | 8,70 | 8,71 |
| 72 | 8,56 | 8,68 |
| 72 | 8,32 | 8,40 |
| 72 | 8,73 | 8,74 |
| 72 | 8,41 | 8,05 |
| 72 | 8,73 | 8,68 |
| 72 | 9,16 | 8,92 |
| 72 | 8,53 | 8,74 |
| 72 | 8,68 | 8,35 |
| 120 | 8,40 | 8,48 |
| 120 | 8,88 | 8,84 |
| 120 | 8,72 | 8,55 |
| 120 | 8,83 | 8,68 |
| 120 | 8,78 | 8,41 |
| 120 | 8,63 | 8,73 |
| 120 | 8,40 | 8,38 |
| 120 | 8,82 | 8,98 |
| 120 | 8,25 | 8,27 |
| 120 | 8,64 | 8,94 |
| 168 | 8,42 | 8,28 |
| 168 | 8,72 | 8,82 |
| 168 | 8,64 | 8,99 |
| 168 | 8,77 | 8,67 |
| 168 | 9,02 | 9,14 |
| 168 | 8,66 | 8,95 |
| 168 | 8,53 | 8,99 |
| 168 | 8,71 | 8,62 |
| 168 | 8,87 | 8,74 |
| 168 | 8,65 | 8,61 |
| 240 | 8,67 | 8,68 |
| 240 | 9,17 | 8,81 |
| 240 | 9,24 | 9,12 |
| 240 | 8,88 | 8,86 |
| 240 | 8,23 | 8,23 |
| 240 | 8,78 | 8,98 |
| 240 | 9,02 | 8,92 |
| 240 | 8,87 | 9,11 |
| 240 | 8,75 | 8,92 |
| 240 | 8,75 | 9,02 |
| 336 | 8,66 | 8,59 |
| 336 | 9,00 | 9,14 |
| 336 | 8,23 | 8,59 |
| 336 | 8,80 | 8,76 |
| 336 | 8,81 | 8,71 |
| 336 | 8,36 | 8,59 |
| 336 | 8,82 | 8,52 |
| 336 | 8,62 | 8,64 |
| 336 | 8,30 | 8,54 |
| 336 | 8,36 | 8,18 |

GF wildtype/A0 coinfection median

| **Time (hours post infection)** | **median of log10 CFU/g feces of Ye wildtype in lumen** | **median of log10 CFU/g feces of Ye YadA0 in lumen** |
| --- | --- | --- |
| 24 | 8,56 | 8,64 |
| 48 | 8,74 | 8,78 |
| 72 | 8,62 | 8,70 |
| 120 | 8,68 | 8,62 |
| 168 | 8,69 | 8,78 |
| 240 | 8,83 | 8,92 |
| 336 | 8,64 | 8,59 |

GF wildtype/T3S0 coinfection

| **Time (hours post infection)** | **log10 CFU/g feces of Ye wildtype in lumen** | **log10 CFU/g feces of Ye T3S0 in lumen** |
| --- | --- | --- |
| 24 | 8,91 | 8,39 |
| 24 | 8,35 | 8,45 |
| 24 | 8,81 | 8,71 |
| 24 | 8,99 | 8,66 |
| 24 | 8,98 | 8,77 |
| 24 | 9,28 | 9,05 |
| 24 | 9,12 | 8,92 |
| 24 | 8,53 | 8,23 |
| 24 | 8,37 | 8,21 |
| 24 | 8,96 | 8,66 |
| 48 | 9,46 | 8,51 |
| 48 | 8,89 | 8,82 |
| 48 | 9,27 | 9,02 |
| 48 | 9,10 | 8,88 |
| 48 | 9,38 | 8,87 |
| 48 | 9,13 | 8,39 |
| 48 | 8,85 | 8,40 |
| 48 | 9,26 | 9,02 |
| 48 | 9,19 | 8,55 |
| 48 | 9,19 | 9,01 |
| 72 | 8,09 | 7,70 |
| 72 | 8,83 | 8,33 |
| 72 | 8,87 | 8,59 |
| 72 | 9,25 | 8,81 |
| 72 | 8,89 | 8,52 |
| 72 | 8,81 | 8,55 |
| 72 | 9,12 | 8,70 |
| 72 | 8,86 | 8,78 |
| 72 | 8,63 | 8,58 |
| 72 | 9,09 | 8,81 |
| 120 | 8,83 | 8,39 |
| 120 | 9,01 | 8,75 |
| 120 | 8,94 | 8,59 |
| 120 | 8,79 | 8,48 |
| 120 | 8,96 | 8,70 |
| 120 | 9,04 | 8,97 |
| 120 | 9,20 | 8,83 |
| 120 | 9,44 | 9,14 |
| 120 | 8,94 | 8,57 |
| 120 | 9,11 | 8,65 |
| 168 | 9,06 | 8,69 |
| 168 | 9,11 | 8,77 |
| 168 | 8,95 | 8,42 |
| 168 | 9,01 | 8,71 |
| 168 | 9,02 | 8,53 |
| 168 | 8,83 | 8,10 |
| 168 | 8,99 | 8,59 |
| 168 | 8,89 | 8,46 |
| 168 | 8,90 | 8,49 |
| 240 | 9,23 | 8,53 |
| 240 | 8,90 | 8,51 |
| 240 | 8,25 | 8,04 |
| 240 | 8,88 | 8,65 |
| 240 | 8,18 | 7,95 |
| 240 | 8,19 | 7,72 |
| 240 | 8,32 | 8,02 |
| 240 | 8,37 | 8,09 |
| 240 | 8,23 | 7,88 |
| 336 | 9,38 | 8,76 |
| 336 | 8,62 | 8,49 |
| 336 | 8,20 | 7,79 |
| 336 | 8,88 | 8,38 |
| 336 | 7,73 | 7,21 |
| 336 | 8,34 | 8,11 |
| 336 | 7,95 | 7,83 |
| 336 | 8,31 | 7,83 |
| 336 | 8,06 | 7,48 |

GF wildtype/T3S0 coinfection median

| **Time (hours post infection)** | **median of log10 CFU/g feces of Ye wildtype in lumen** | **median of log10 CFU/g feces of Ye T3S0 in lumen** |
| --- | --- | --- |
| 24 | 8,94 | 8,66 |
| 48 | 9,19 | 8,85 |
| 72 | 8,87 | 8,59 |
| 120 | 8,99 | 8,68 |
| 168 | 8,99 | 8,53 |
| 240 | 8,32 | 8,04 |
| 336 | 8,31 | 7,83 |

MyD88^-/-^ wildtype/A0 coinfection

| **Time (hours post infection)** | **log10 CFU/g feces of Ye wildtype in lumen** | **log10 CFU/g feces of Ye YadA0 in lumen** |
| --- | --- | --- |
| 16 | 4,90 | 4,18 |
| 16 | 5,26 | 4,06 |
| 16 | 4,76 | 3,65 |
| 16 | 4,50 | 4,29 |
| 16 | 4,71 | 4,38 |
| 16 | 1,79 | 2,07 |
| 16 | 4,14 | 3,90 |
| 16 | 3,88 | 2,76 |
| 16 | 5,03 | 4,63 |
| 16 | 3,45 | 2,81 |
| 24 | 2,64 | 2,34 |
| 24 | 3,17 | 3,18 |
| 24 | 2,74 | 1,74 |
| 24 | 3,62 | 3,13 |
| 24 | 3,76 | 2,05 |
| 24 | 3,77 | 2,24 |
| 24 | 4,14 | 2,27 |
| 24 | 3,06 | 2,46 |
| 24 | 5,23 | 3,23 |
| 24 | 3,97 | 3,27 |
| 40 | 3,55 | 4,17 |
| 40 | 5,25 | 5,73 |
| 40 | 3,18 | 1,79 |
| 40 | 5,63 | 4,25 |
| 40 | 4,62 | 1,63 |
| 40 | 5,63 | 1,79 |
| 40 | 4,32 | 2,28 |
| 40 | 3,90 | 3,14 |
| 40 | 5,54 | 5,35 |
| 40 | 3,27 | 2,42 |
| 48 | 3,74 | 4,84 |
| 48 | 5,27 | 5,68 |
| 48 | 5,67 | 1,79 |
| 48 | 6,33 | 3,62 |
| 48 | 6,01 | 1,79 |
| 48 | 6,27 | 1,53 |
| 48 | 4,51 | 1,83 |
| 48 | 3,34 | 2,63 |
| 48 | 4,73 | 6,55 |
| 48 | 3,10 | 2,12 |

MyD88^-/-^ wildtype/A0 coinfection median

| **Time (hours post infection)** | **median of log10 CFU/g feces of Ye wildtype in lumen** | **median of log10 CFU/g feces of Ye YadA0 in lumen** |
| --- | --- | --- |
| 16 | 4,61 | 3,98 |
| 24 | 3,69 | 2,40 |
| 40 | 4,47 | 2,78 |
| 48 | 5,00 | 2,38 |

MyD88^-/-^ wildtype/T3S0 coinfection

| **Time (hours post infection)** | **log10 CFU/g feces of Ye wildtype in lumen** | **log10 CFU/g feces of Ye T3S0 in lumen** |
| --- | --- | --- |
| 16 | 3,45 | 3,20 |
| 16 | 3,46 | 3,39 |
| 16 | 3,16 | 4,03 |
| 16 | 4,52 | 3,81 |
| 16 | 2,65 | 2,65 |
| 16 | 3,23 | 3,58 |
| 16 | 3,61 | 3,17 |
| 16 | 3,99 | 3,42 |
| 16 | 3,17 | 3,58 |
| 16 | 3,42 | 3,79 |
| 24 | 2,24 | 2,24 |
| 24 | 2,80 | 4,48 |
| 24 | 5,98 | 4,38 |
| 24 | 2,24 | 2,24 |
| 24 | 1,94 | 1,94 |
| 24 | 3,42 | 2,24 |
| 24 | 5,11 | 3,04 |
| 24 | 3,07 | 2,04 |
| 24 | 3,65 | 2,51 |
| 24 | 2,24 | 2,24 |
| 40 | 2,71 | 3,19 |
| 40 | 4,59 | 5,52 |
| 40 | 6,52 | 5,66 |
| 40 | 4,57 | 3,57 |
| 40 | 5,32 | 3,20 |
| 40 | 4,63 | 3,58 |
| 40 | 5,94 | 4,59 |
| 40 | 3,76 | 3,31 |
| 40 | 3,86 | 3,00 |
| 40 | 4,25 | 3,56 |
| 48 | 2,96 | 3,36 |
| 48 | 3,88 | 3,65 |
| 48 | 6,61 | 5,20 |
| 48 | 5,34 | 2,24 |
| 48 | 5,21 | 3,20 |
| 48 | 4,25 | 4,03 |
| 48 | 4,72 | 4,34 |
| 48 | 3,24 | 3,16 |
| 48 | 3,55 | 1,95 |
| 48 | 3,64 | 3,19 |

MyD88^-/-^ wildtype/T3S0 coinfection median

| **Time (hours post infection)** | **median of log10 CFU/g feces of Ye wildtype in lumen** | **median of log10 CFU/g feces of Ye T3S0 in lumen** |
| --- | --- | --- |
| 16 | 3,44 | 3,50 |
| 24 | 2,94 | 2,24 |
| 40 | 4,58 | 3,57 |
| 48 | 4,07 | 3,28 |

CFU values at detection limit are labeled.
