## Supplemental File 15 MatLab Script for "Model-based prediction of bacterial population dynamics in gastrointestinal infection"

```
function paramfit7D
% main program for fitting parameters of an ODE model to data(SPF)
% the model and the error function are defined in the file SfunD.m
clearvars -global
global tdata xdata ydata x0 b c d

%% reading the data from the excel file
tdata=xlsread('Yersinia_ExperimentalData.xlsx','SPFA0','A:A')
xdata=xlsread('Yersinia_ExperimentalData.xlsx','SPFA0','B:B')
ydata=xlsread('Yersinia_ExperimentalData.xlsx','SPFA0','C:C')

%% initial condition
x0(1) = 1000+rand(1,1)*(100000-1000);
x0(2) = 10;
x0(3) = 10;
x0(4) = 1000000+rand(1,1)*(1000000000-1000000);
x0(5) = 1000;
x0(6) = 1000;
x0(7) = 0;
c(1) = 1;                               % Maximum immunity action
c(2) = 1/4;                             % Rate at which intestines are
discharged
c(3) = 1;                               % Maximum capacity of the immune
system
d(1) = 3/2.3;                           % Kottel factor


%% initial guess of parameter values
b(1) = 0.4+rand(1,1)*(2-0.4);           % Max growth rate of intestinal
bacteria
b(2) = 0.4+rand(1,1)*(2-0.4);           % Max growth rate of wild-type
Yersinia which is supposed to be equal with Max growth rate of mutant
Yersinia
b(3) = b(2);                            % Max growth rate of mutant Yersinia
b(4) = x0(1);                           % Carrying capacity of the mucosa
b(5) = x0(4);                           % Carrying capacity of the lumen
b(6) = 0.001+rand(1,1)*(0.11-0.001);    % Immunity adjustment factor for
wild-type Yersinia
b(7) = 0.11+rand(1,1)*(0.2-0.11);       % Immunity adjustment factor for
mutant Yersinia
b(8) = 0.0004+rand(1,1)*(1-0.0004);     % Max. rate of immune growth
b(9) = 0.5+rand(1,1)*(1-0.5);           % Standard deviation
fprintf('%i\n', b)

%% minimization step
    options=optimset('TolFun',1e-7,'TolX',1e-7,'Display','iter',
'MaxFunEvals',150);
    [bmin, Smin] = fminsearchbnd(@Sfun7D,d,[0.4 0.4 0.4 1000 1000000 0.001
0.11 0.0004 0.5], [2 2 2 10000000 1000000000 0.11 0.2 1 1], options);
    disp('Estimated parameters b(i):');
    disp(bmin)
    disp('Smallest value of the error S:');
    disp(Smin)
end
function S = Sfun7D(b)
```

```
% computation of an error function for an ODE model
% INPUT: b - vector of parameters
global tdata xdata ydata x0 b c d

%% ODE model
% (nested function, uses parameters b(i) of the main function)
%% dx(1) = Commensal bacteria in mucosa
%% dx(2) = wild-type yersinia in mucosa
%% dx(3) = mutant yersinia in mucosa
%% dx(4) = Commensal bacteria in lumen
%% dx(5) = wild-type yersinia in lumen
%% dx(6) = mutant yersinia in lumen
%% dx(7) = Immune system
function dx = f(t,x)
         dx = zeros(7,1);
         dx(1) = b(1)*(1-((x(1)+x(2)+x(3))/b(4)))*x(1) - c(1)*x(7)*x(1);
         dx(2) = b(2)*(1-((x(1)+x(2)+x(3))/b(4)))*x(2) -
c(1)*b(6)*x(7)*x(2);
         dx(3) = b(3)*(1-((x(1)+x(2)+x(3))/b(4)))*x(3) -
c(1)*b(7)*x(7)*x(3);
         dx(4) = b(1)*(1-((x(4)+x(5)+x(6))/b(5)))*x(4) - c(2)* x(4);
         dx(5) = b(2)*(1-((x(4)+x(5)+x(6))/b(5)))*x(5) - c(2)* x(5) +
b(2)*((x(1)+x(2)+x(3))/b(4))*x(2);
         dx(6) = b(3)*(1-((x(4)+x(5)+x(6))/b(5)))*x(6) - c(2)* x(6) +
b(3)*((x(1)+x(2)+x(3))/b(4))*x(3);
         dx(7) = b(8)*(x(2)+x(3))*(c(3)-x(7));
        end
%% numerical integration set up
tspan = [0:1:max(tdata)];
[tsol,xsol] = ode45(@f,tspan,x0);

%% find predicted values x(tdata)
xpred = interp1(tsol,xsol(:,5),tdata);
ypred = interp1(tsol,xsol(:,6),tdata);

%% compute total error with Maximum Likelihood Method
S = 0; S1 = 0; S2 = 0; S3 =0 ; S4 = 0; Sm1 = 0; Sm2 = 0; Sm3 = 0; Sm4 = 0;
for i = 1:length(tdata)
    if xdata(i,1) <= 2.06 && ydata(i,1) <= 2.06
            g = @(x) exp(-(x-log10(d(1)*xpred(i,1))).^2/(2*b(9)^2));
            g1 = integral(g,-inf,2.05);
            g2 = log(b(9)) - log(g1);
            S1 = S1 + g2;
            h = @(x) exp(-(x-log10(d(1)*ypred(i,1))).^2/(2*b(9)^2));
            h1 = integral(h,-inf,2.05);
            h2 = log(b(9)) - log(h1);
            S3 = S3 + h2;
            Sm1 = Sm1 + S1 + S3;
    elseif xdata(i,1) > 2.06 && ydata(i,1) > 2.06
            S2 = S2 + ((log10(d(1)*xpred(i,1))-xdata(i,1)).^2/(2*(b(9)^2)))
+ log(b(9));
            S4 = S4 + ((log10(d(1)*ypred(i,1))-ydata(i,1)).^2/(2*(b(9)^2)))
+ log(b(9));
            Sm2 = Sm2 + S2 + S4;
    elseif xdata(i,1) <= 2.06 && ydata(i,1) > 2.06
            g = @(x) exp(-(x-log10(d(1)*xpred(i,1))).^2/(2*b(9)^2));
```

```
            g1 = integral(g,-inf,2.05);
            g2 = log(b(9)) - log(g1);
            S1 = S1 + g2;
            S4 = S4 + ((log10(d(1)*ypred(i,1))-ydata(i,1)).^2/(2*(b(9)^2)))
+ log(b(9));
            Sm3 = Sm3 + S1 + S4;
    else
            S2 = S2 + ((log10(d(1)*xpred(i,1))-xdata(i,1)).^2/(2*(b(9)^2)))
+ log(b(9));
            h = @(x) exp(-(x-log10(d(1)*ypred(i,1))).^2/(2*b(9)^2));
            h1 = integral(h,-inf,2.05);
            h2 = log(b(9)) - log(h1);
            S3 = S3 + h2;
            Sm4 = Sm4 + S2 + S3;
    end
        S = S + Sm1 + Sm2 + Sm3 + Sm4;
end
end
```
